# Selective refueling of CAR T cells using ADA1 and CD26 boosts antitumor immunity

**DOI:** 10.1101/2023.09.18.557956

**Authors:** Yue Hu, Abhijit Sarkar, Kevin Song, Sara Michael, Magnus Hook, Ruoning Wang, Andras Heczey, Xiaotong Song

## Abstract

CAR T cell therapy has revolutionized the treatment of hematologic malignancies, but its efficacy in solid tumors remains limited due to the immunosuppressive nature of the tumor microenvironment and the inability of T cells to persist and traffic to the tumor site. While current strategies focus on enhancing CAR T cell activity through costimulatory molecules and cytokines, a critical yet often overlooked factor is the competition for nutrients between tumor cells and T cells in the nutrient-deprived tumor microenvironment. To address this challenge, we employed a selective metabolic refueling (MR) strategy by providing T cells with inosine as an alternative fuel source for growth and functionality. In this study, we engineered CAR T cells to co-express a membrane-bound CD26 and a cytoplasmic adenosine deaminase 1 (ADA1) fused to an anti-CD3 scFv. ADA1 irreversibly converts both intracellular and extracellular adenosine to inosine, overcoming adenosine-mediated immunosuppression and providing T cells with inosine for growth. The inclusion of an anti-CD3 scFv fusion partner and overexpressing CD26 boosts ADA1 capture in a membrane proximal manner, providing inosine for T cells and minimizing feeding the tumor cells. We demonstrate that ADA1 is conditionally secreted only in stress conditions and that it activates CAR T cells through trans-signaling in a tumor-specific manner. In addition, we show that, compared to unmodified CAR T cells, CD26-overexpressing CAR T cells have better migration capacity and are less susceptible to TGF-β1 suppression. Finally, we found that, in mice models of human hepatocellular carcinoma (GPC3-MR-CAR) and human non-small cell lung cancer (HER2-MR-CAR), metabolically refueled CAR T cells (MR-CAR) are more efficient in reducing tumor growth than unmodified CAR T cells. Thus, selective refueling CAR T cells using ADA1 and CD26 holds promise for improving the efficacy of CAR T cell therapy of solid tumors.

## Introduction

Chimeric antigen receptor (CAR) T cell therapy is a promising approach that combines the specificity of monoclonal antibodies with the targeted biodistribution and long-term persistence of effector lymphocytes to selectively target and eliminate malignant cells. Early and late-phase clinical trials have shown breakthrough successes in patients with hematologic malignancies treated with CAR T cells, leading to six FDA-approved CAR T cell therapies^1–6^. However, CAR T cell therapy has shown only modest results in patients with solid tumors. The limited efficacy is likely multicausal including limited CAR T cell trafficking to solid tumors, tumor-mediated immunosuppression, and limited expansion and persistence of CAR T cells in the tumor microenvironment (TME)^7,8^. Current strategies primarily use co-stimulatory molecules and more recently cytokines to optimize CAR T cell activation and survival^9–11^. However, nutritional competition between tumor cells and T cells poses a significant but underappreciated challenge to the efficacy of CAR T cells in solid tumors^12–18^. Because cancer cells have high metabolic demands, the TME creates a metabolic stress environment that can affect the function of CAR T cells by limiting energy resources like glucose and by producing immunosuppressive metabolites like adenosine. The majority of glucose is absorbed by tumor cells, leaving CAR T cells starved and unable to survive regardless of stimulations.

Our previous research has shown that inosine can act as a substitute carbon source to support T cell proliferation and activity in the absence of glucose^19^. T cells can metabolize inosine to hypoxanthine and phosphorylated ribose through purine nucleoside phosphorylase^19,20^. The ribose component of inosine can enter into central metabolic pathways to provide energy in the form of ATP and to support biosynthetic processes. In addition, inosine can modulate immune responses through the activation of A2AR and A3R in context-dependent manners. Notably, in the presence of IFN-γ, inosine significantly boosted Th1 differentiation of naïve T cells, whereas in the absence of IFN-γ, inosine inhibited this pathway^21^. Given the ability of adoptive T cell transfer and immune checkpoint inhibitors to convert a suppressive TME into a supportive one, it is reasonable to speculate that inosine can increase the effectiveness of T cell therapy or immune checkpoint inhibitors in combating tumors. Indeed, inosine has been demonstrated to enhance tumor immunogenicity, making tumor cells more vulnerable to the cytolytic effects of immune cells^22^. However, as inosine can also act as a fuel for tumor growth, targeted delivery of inosine to CAR T cells is necessary to optimize CAR T cell therapy.

Inosine is a nucleoside produced from adenosine via Adenosine deaminase (ADA)^23^. There are two isoenzymes of ADA present in humans, ADA1 and ADA2^24–26^. ADA1 is primarily a cytoplasmic protein that is ubiquitously expressed by most body cells^23,25,27^, but although ADA1 lacks a signal peptide, it has been shown to be secreted via a non-classic pathway and ADA1 is found in human plasma^26,28,29^. ADA2 is the predominant isoform found in human plasma. ADA1 and ADA2 play different parts in regulating immune responses, independent of their ADA activity. ADA1 has co-stimulatory effects on T-cell-mediated immunity by engaging T cells that express the ADA1 receptor CD26^28,30,31^, whereas ADA2 in a CD26 independent manner, binds to immune cells, including neutrophils, CD16+ monocytes, NK cells, B cells, and regulatory T cells^32,33^. Both ADA1 and ADA2 are significantly increased in multiple human tumor tissues^25^. However, ADA1 expression positively correlates with the expression of genes associated with cell division and exhibit only a moderate positive correlation with genes associated with tumor-infiltrating lymphocytes. In contrast, ADA2-expression correlates with genes associated with immune responses in multiple types of infiltrating immune cells. These observations suggest that ADA1’s primary function is in adenosine metabolism, while ADA2 plays a more significant role in regulating immune responses. This functional disparity is aligned with the observation that ADA1 has a 100-fold higher affinity for adenosine compared to ADA2. Notably, high expression levels of ADA1 in tumor tissues have been associated with worse outcomes, while high ADA2 expression levels are linked to favorable results in a variety of cancers^25^. This raises questions about whether enhanced ADA1 activity broadly in tumor tissues provides an additional energy source for the growth of tumor cells. Therefore, even though CAR T cells overexpressing secreted ADA1 have shown the potential to significantly slow tumor growth in mouse models^34^, it may be crucial to prevent inosine from becoming an extra energy source for cancer cells in the treatment of human cancer.

The interaction of ADA1 with T cells through CD26 is an important step in the co-stimulation and activation of T cells. After T cells are activated, the multifunctional protein CD26 is increased on their plasma membranes, and its expression is closely controlled during T cell development^30,31,35,36^. Upon activation, CD26 recruits CARMA1 to its cytosolic domain, leading to NF-kB activation, T cell proliferation, and IL-2 production. CD26-mediated co-stimulation differs from CD28 co-stimulation and preferentially results in cytotoxicity through production of granzyme B, tumor necrosis factor-α, interferon-γ, and Fas ligand^37^. Additionally, CD26^high^ T cells have been shown to elicit tumor immunity against various cancers, with enhanced migration and persistence, chemokine receptor profile, cytotoxicity, resistance to apoptosis, and stemness^38^. However, TGF-β1 can significantly downregulate CD26 expression and impair the cell activities, emphasizing the necessity to overexpress CD26 in CAR T cells to preserve their functionality^39,40^. In this study, we engineered CAR T cells with a helper vector to co-express CD26 and ADA1.CD3scFv, which facilitated the capture of ADA1 in a membrane-proximal manner. We found that this combination selectively enhanced the proliferation and cytokine production of CAR T cells, without affecting tumor cells, resulting in increased cytolytic activity against cancer cells. Furthermore, co-expression of CD26 and ADA1.CD3scFv improved the persistence of CAR T cells in the tumor microenvironment and increased their migration towards the tumor, resulting in better tumor control. Our study highlights the potential of combining CD26 and ADA1 in CAR T cell engineering as a promising strategy to improve the efficacy of CAR T cell therapy.

## Methods

### Cell lines

The cell lines 293T, A549, Calu3, and HepG2 were obtained from the American Type Culture Collection. The cell line Huh-7 was a kind gift from Dr. Andras Heczey (Texas Children’s Hospital, Baylor College of Medicine, Houston, TX). These cell lines were cultured in Dulbecco’s modified Eagle media with 10% fetal bovine serum (FBS). Jurkat-Lucia NFAT Cells (Invivogen, jktl-nfat) and Jurkat-Dual Cells (Invivogen, jktd-isnf) were used as reporter cell lines and maintained as per the instructions provided.

### Generation of retroviral constructs

The Moloney murine leukemia virus–derived SFG retroviral vector backbone, which has been clinically validated, was generously provided by Dr. Andras Heczey of Baylor College of Medicine ^41,42^. The retroviral vector was used to encode four distinct ADA1 constructs, including cytoplasmic ADA1, secreted ADA1, permanent integral membrane-bound ADA1, or ADA1 fusion protein with anti-human CD3 scFv (ADA1.CD3scFv). The genes were synthesized by Genscript and subcloned into SFG gamma-retroviral vectors. To express integral membrane-bound ADA1, a human CD28 transmembrane domain was added to the C-terminal of ADA1. To express ADA1.CD3scFv, the ADA1 gene was fused with an anti-human CD3 scFv and was separated by a (G4S)3 linker. The anti-human CD3 scFv was obtained from the OKT3 clone.

To generate ADA1.CD3scFv/CD26 vector (MR vector), codon-optimized genes encoding CD26, T2A, and ADA1.CD3scFv were synthesized by Genscript and subcloned into SFG gamma-retroviral vectors. To generate HER2-CAR, codon-optimized genes encoding anti-human HER2 scFv derived from the FRP5 clone, CD28 transmembrane domain, CD28 endodomian, and CD3ζ were synthesized by Genscript and subcloned into SFG gamma-retroviral vector. This HER2-specific CAR is identical to the second-generation HER2 CAR, which has been elevated in several clinical studies ^43–45^. The GPC3-CAR retroviral vector encoding anti-human GPC3 scFv derived from the GC33 clone, CD28 transmembrane domain, 4-1BB endodomain, and CD3ζ was also generously provided by Dr. Andras Heczey from Baylor College of Medicine^41,42^.

### Retrovirus production and transduction of primary T cells

To generate CAR T cells, retroviral supernatants were produced by transient transfection of 293T cells with retroviral vector-containing plasmids of GPC3-CAR construct, HER2-CAR construct, or 50-50 mixtures of MR vector with CAR constructs. PegPam3 plasmid encoding the MoMLV gag-pol was used to generate retroviral supernatants as previously described^41,42^. Human peripheral blood mononuclear cells purchased from STEMCELL (70025) was stimulated by OKT-3 (Miltenyi Biotec, 130-093-387) and CD28 (BD Pharmingen, 567117) mAb-coated plates. The cells were washed and replated in complete RPMI 1640 medium with 100 U/ml IL-2 (PeproTech, 200-02) 48 hours later. The cells were then maintained in incubators at 37°C with 5% CO2 for expansion.

### Western blot

Cells were washed with PBS and lysed using cell lysis buffer containing Protease and Phosphatase Inhibitor Cocktail (ThermoFisher; 78440). 30 μg of quantified proteins were separated on Bis-Tris pre-cast SDS-PAGE mini-gel (ThermoFisher; NP0322) and transferred to polyvinylidene fluoride membrane using a dry blotting system (ThermoFisher). After blocking the membranes were incubated overnight with desired ADA1 primary antibody (Sigma Aldrich, HPA001399) and then incubate 1 hour with secondary antibody. Protein bands were visualized using a ChemiDoc XRS imager system (Bio-Rad).

### qPCR

HER2-CAR, HER2-MRCAR, GPC3-CAR, and CP3-MRCAR was cultured, and mRNA was gathered using RNeasy Plus mini kit (Qiagen #74134) according to the manufacturer’s instruction for qPCR. Briefly, 1000ng of RNA was used as a template for cDNA synthesis in 10µL reaction volume (Bio-Rad #1725038). qPCR was performed using Luna universal QPCR mix (NEB # M3003) in the CFX96 Real-Time PCR system with a C1000 Thermal Cycler (Bio-Rad). Results are represented as fold change above control after normalization to GAPDH. IFN gamma forward primer: GAGTGTGGAGACCATCAAGGA; Reverse primer: TGTATTGCTTTGCGTTGGAC.

### Measure of cytokines

To measure IFN gamma concentrations, the Human IFN gamma ELISA kit (R&D Systems, DIF50C) was used according to the manufacturer’s instructions. 1×105, 5×104, and 2.5×104 HER2-CAR or HER2-MR-CAR cells were cultured in presence or absence of A549 at a 1:1 ratio. Cell Culture supernatants were collected at 24 hours, centrifuged, and frozen until the assay.

### LDH assay

Cytotoxicity of GPC3-CAR T cells and HER2-CAR T cells were assessed and compared to GPC3-MR-CAR T cells and HER2-MR-CAR T cells using LDH assay. The GPC3-CAR and GPC3-MR-CAR T cells were co-cultured with HepG2 or Huh7 cells, and the HER2-CAR and HER2-MR-CAR T cells were co-cultured with A549 or Calu3 cells at varying E:T ratios. Cell culture supernatants were collected and measured using LDH Cytotoxicity WST Assay (ENZO, ENZ-KIT157) according to the manufacturer’s instructions.

### ADA activity determination

Cells were cultured at indicated cell density for 24 hours. Then cell culture supernatants and cells were collected for determination of ADA activity. ADA activity was measured using Adenosine Deaminase Assay Kit (GenWay Biotech, GWB-BQK080) according to the manufacturer’s instructions.

### Inosine assay

Supernatant of equal amount of tumor tissue homogenization from Non-transduced, HER2-CAR and HER2-MR-CAR group from ex vivo analysis was used to perform Inosine assay according to Cell BioLabs Kit (MET5092) protocol and the inosine concentration was calculated from the standard curve.

### Migration assay

Transwell inserts (Falcon, 0.4um) were used in 24 well tissue culture plate. 4×10^5^ Non transduced, HER2-CAR and HER2-MRCAR cells were placed in the top well of a transwell plate (Cell Migration/Chemotaxis Assay Kit, ab235696). 235 ul of A549 cancer cells supernatant was placed in bottom well as chemoattractant. Supernatant from A549 cancer cells were collected 24 hours after plating. The ability of cells to migrate assayed by both fluorescence activity and cell counting as per KIT protocol.

### RNA sequencing

RNA-seq was performed in triplicate for each experimental group. Six days after transduction RNA was isolated using the RNeasy Plus mini kit (Qiagen). RNA was checked for quality control and then sequenced by Genewiz form Azenta Life Sciences. After getting the Fastq file reads were mapped to the human reference genome (hg38) using STAR (v2.7.2b) RNAseq alignment tool. Transcript levels were quantified to the reference genome using a Bayesian approach. Normalization was done using counts per million (CPM) method. Differential expression was done using DESeq2 (v3.5) with default parameters. The normalized counts of each gene were log2 scaled and ranked by their fold changes values between paired samples. A ranked list of genes with log2[FC] values was used in pre-ranked GSEA analysis. Hallmark genesets were utilized as background genesets database in GSEA. Broad institute’s standalone GSEA (version 4.0.3) software was used to perform enrichment analysis.

### Cell sorting

Tumor tissue were collected from mice and filtered through 70 um cell strainers (CELLTREAT Scientific Products, 229483) for single-cell suspensions. The filtered cells were then stained with CD3-PE (BioLegend, 300407) for Flow cytometry sorting (BD FACSAria Fusion).

### Flow Cytometry

Anti-F(ab)_2_ Alexa Fluor 647-conjugated antibody (Jackson ImmunoResearch, 115-605-006) and anti-goat IgG1 isotype control (Jackson ImmunoResearch, 115-605-006) is used to detect GPC3-CAR expression. Alexa flour 647 conjugated HER2 protein (Biotechne, AFR1129) was used to detect HER2 expression using a similar method as listed above. The antibodies used for T-cell phenotyping and cytokine production analyses are: CD3-BV785 (BioLegend, 317329), CD8-Pacific Blue (BioLegend, 344717), CCR2-BUV395 (BD, 747854), CCR5-FITC (BioLegend, 313705), CD26-FITC (BD, 555436), ADA1-PE (Santa Cruz, sc-28346), PD-1-Alexa flour 647 (BD, 566851), IL2-BV711 (BD #563946), IFN gama-PE-CF594 (BD, 562392), GZMB-APC (BioLegend, 372203), Perforin-BV711 (BioLegend, 08129), T-bet-BV785 (BioLegend, 644835), IFN-γ-BV570 (BioLegend, 502534), GZMB-PE (BioLegend, 372208) and IL-2-BV711 (BD, 563946). Intracellular cytokine staining was performed using a Transcription Factor Buffer Set (BD Biosciences, 562574). Flow cytometry assessment was performed on ZE5 Cell Analyzer (BIO-RAD). Results were analyzed with FlowJo software (TreeStar).

### CFSE assay

Cells were stained with Carboxyfluorescein diacetate succinimidyl ester (CFSE) Cell Division Tracking Kit (BioLegend, 423801), and then cultured in complete RPMI1640 with IL-2 for 3 days. On day 3, cells were harvested and the CFSE fluorescent dilution was analyzed by flow cytometry.

### In Vivo experiments

6–8-week-old NOD.Cg-PrkdcSCID Il2rgtm1Wjl/SzJ (NSG) mice were purchased from The Jackson Laboratory and maintained at the PAR Facility of Texas A&M Institute of Biosciences and Technology and all animal experiments were approved by the Institutional Animal Care and Use Committee of Texas A&M University College of Medicine, Institute of Biosciences and Technology. Tumor cells or CAR T cells were diluted in 100 μl normal saline and were injected via indicated routes. Briefly, 6-to 10-week-old NSG mice were injected subcutaneously with different tumor cell lines respectively to generate tumor xenograft models. Tumor volume was determined by caliper measurement (LxW2/2). Once tumors reached 4–6 mm in diameter, indicated number of CAR T cells and MR-CAR T cells were injected intravenously (i.v.) via tail vein. Mice were assessed daily, and tumor sizes were measured three times a week. For ex vivo analysis of tumor infiltrating lymphocytes, CAR T cells were injected when tumors reached an average size of 200–300 mm^3^. At the indicated time points, tumors were dissected, and tumor infiltrating lymphocytes were analyzed.

### Statistical analyses

Descriptive analysis was performed to summarize data for cytokine production and CAR expression on T cells. Response variables were transformed, if necessary, to achieve normality. Unpaired two-tailed Student’s t test was used to assess differences between two experimental groups. Graphs that incorporate error bars present the mean as the central value, while the error bars represent the standard error of the mean (SEM). A p value <0.05 was considered statistically significant. Since these experiments were exploratory in nature, there was no estimation to determine the appropriate sample size. Therefore, we relied on traditional sample sizes of greater than 5 for our animal studies.

## Results

### Rational design of a helper vector optimizing ADA1 function in CAR T cells

ADA1 overexpressed in CAR T cells recently was shown to have some positive effects on reducing tumor size in mice^34^. However, the high expression of ADA1 seen in human cancers and its link to poor survival outcomes raise questions about whether such treatments could also promote the growth of cancerous cells in humans. Therefore, to harness the benefits of ADA1 in T cell therapy, it is essential to maximize its potential in boosting T cell function, while avoiding inosine to become an energy source for cancer cells. To selectively boost the immunobiology of ADA1 in CAR T-cells, we designed a helper vector to metabolically reprogram (MR) T cells. The MR vector drives the expression of a membrane bound CD26 and an ADA1 protein fused to anti-CD3 scFv in the T cells to support T cell proliferation and activation **(Figure 1A)**. Furthermore, we designed a series of constructs to overexpress secreted ADA1, cytoplasmic ADA1, or membrane-bound ADA1, and the impact of these constructs on modulating T cell function was compared **(Figure 1A)**.

**Figure 1.**
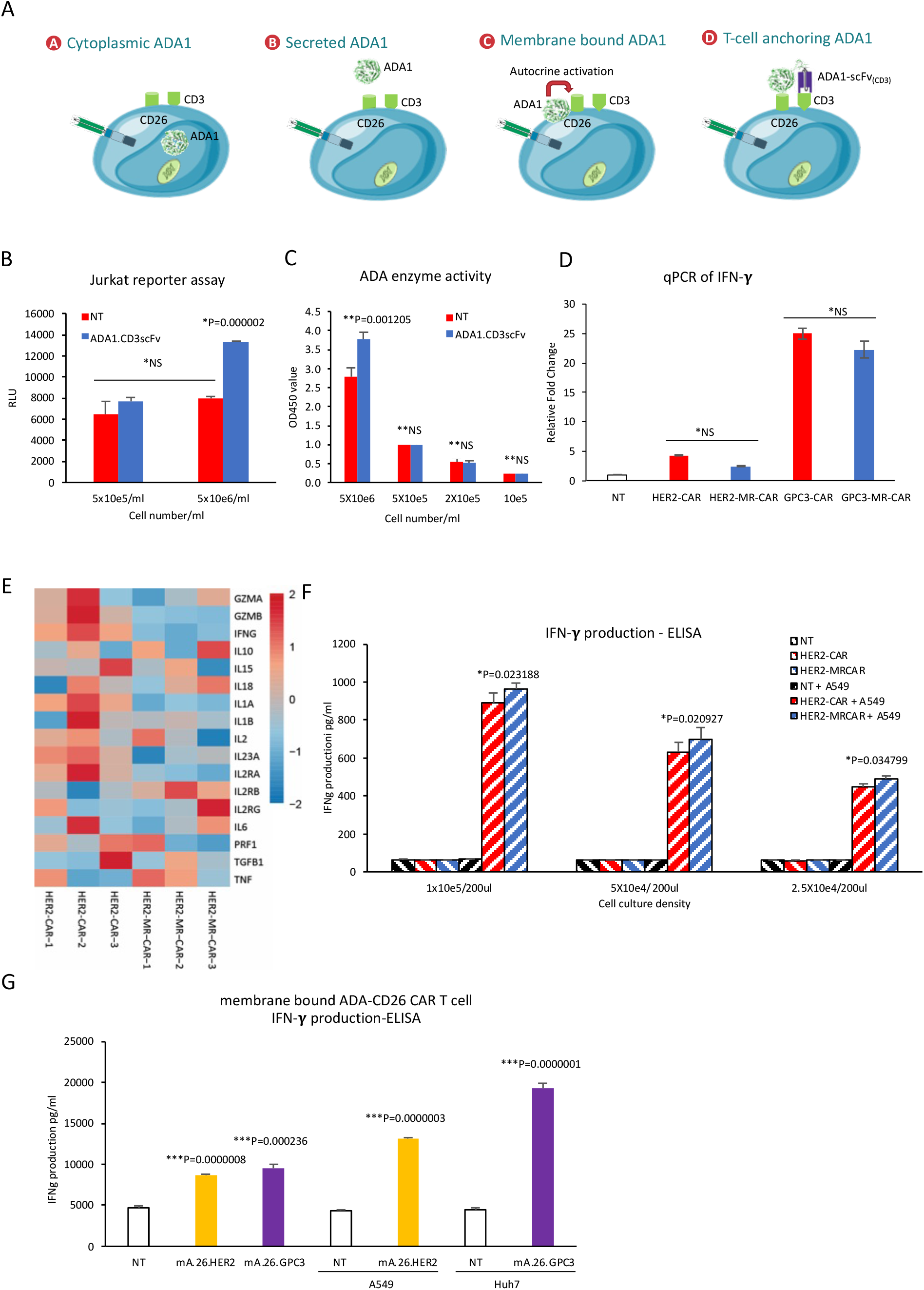
Rational design and implementation of ADA1 in CAR T cells. **(A)** Four different designs of ADA1 were designed for use in CAR T cell therapy. **(B)** The conditional secretion of ADA1.CD3scFv was tested using NFAT luciferase reporter Jurkat T cells. MR vector transduced NFAT luciferase reporter Jurkat T cells (ADA1.CD3scFv) were cultured in either high cell density (5×10^6^ cells per ml, stress condition) or low cell density (5×10^5^ cells per ml, physiological condition) for 24 hours. A reporter assay was then conducted to determine the secretion of ADA1.CD3scFv. Non-transduced NFAT luciferase reporter Jurkat T cells (NT) were used as control. *NS (no significant difference) between NT in physiological condition, ADA1.CD3scFv in physiological condition, and NT in stress condition. **P=0.000002 for ADA1.CD3scFv in stress condition verse NT in stress condition or ADA1.CD3scFv in physiological condition. **(C)** An ADA enzyme activity assay was conducted to determine the conditional secretion of ADA1.CD3scFv. MR vector transduced NFAT luciferase reporter Jurkat T cells (ADA1.CD3scFv) were cultured in either high cell density (5×10^6^ cells per ml, stress condition) or low cell density (5×10^5^ cells per ml, physiological condition) for 24 hours. Cells were collected for ADA activity assay. Non-transduced NFAT luciferase reporter Jurkat T cells (NT) were used as control. *NS for ADA1.CD3scFv in physiological condition verse NT in physiological condition. **P=0.001205 for ADA1.CD3scFv in stress condition verse NT in stress condition. **(D)** qRT-PCR was used to measure the expression of IFN-γ in both HER2-specific and GPC3-specific MR-CAR T cells. HER2-specific or GPC3-specific CAR T cells were cultured without tumor cells for 24 hours. Cells were collected, and mRNA was extracted. IFN-γ levels were then measured by qRT-PCR. *NS for either HER2-MR-CAR verse HER2-CAR or GPC3-MR-CAR verse GPC3-CAR. **(E)** mRNA sequencing was used to analyze inflammatory cytokines, Granzyme A, and Granzyme B in CAR T cells. **(F)** ELISA was used to measure IFN-γ expression in HER2-specific MR-CAR T cells both in the presence and absence of HER2-positive A549 NSCLC. HER2-MR-CAR T cells, HER2-CAR T cells, or NT were cultured at indicated density either with or without A549 tumor cells for 24 hours. After incubation, the culture medium was collected, and IFN-γ levels were measured using ELISA. *NS (no significant difference) for HER2-MR-CAR verse HER2-CAR. *The figure indicated the P values for HER2-MR-CAR+A549 verse HER2-CAR+A549. **(G)** ELISA was used to measure IFN-γ expression in membrane-bound ADA1 and CD26 overexpressed CAR T cells. Human PBMCs were first activated by anti-CD3/CD28 mAbs and then transduced with a retroviral vector encoding both membrane-bound ADA1 and CD26, as well as either HER2-specific CAR (mA.26.HER2) or GPC3-specific CAR (mA.26.GPC3). Non-transduced, anti-CD3/CD28 activated PBMCs were used as a control (NT). Next, mA.26.HER2 or NT were cultured either with or without A549 tumor cells, while mA.26.GPC3 or NT were cultured either with or without Huh7 cells. After 24 hours later, the culture medium was collected, and IFN-γ levels were measured using ELISA. *P=0.0000008 and P=0.000236 for mA.26.HER2 and mA.26.GPC3 verse NT (without tumor cells co-culture). **P=0.0000003 for mA.26.HER2 verse NT (with A549 co-culture). ***P=0.0000001 for mA.26.GPC3 verse NT (with Huh7 co-culture).

To induce the secretion of ADA1 from human T cells, the human IL-2 signal peptide was inserted into the N-terminal of ADA1^46^. The secretion of ADA1 was evaluated through ADA1 ELISA and ADA1 enzyme activity assay in the T cell culture medium. Unexpectedly, the expression of secreted ADA1 in the cell culture medium was not detectable at 24 hours or 48 hours. This observation suggests that the mechanism of ADA1 secretion in human T cells may be different from that of other proteins and may not be activated by the IL-2 signal peptide.

We then asked if cellular stress or activation may cause the secretion of ADA1 in human T cells given that lymphocytes can release ADA1 in response to an acidic environment or stimulation with anti-CD3 antibodies^28,29^. To address this possibility, the secretion of ADA1.CD3scFv was evaluated using NFAT luciferase reporter human Jurkat T cells. NFAT luciferase reporter human Jurkat T cells were transduced with the retroviral vector encoding ADA1.CD3scFv and then cultured at high cell density (5 million cells per ml) or low cell density (0.5 million cells per ml). High cell density should result in cellular stress due to higher lactate concentrations, a lower medium pH, and a lack of nutrients. After 24 hours of culture, the Jurkat T cells were subjected to luciferase assay to determine the CD3scFv-mediated activation of Jurkat T cells **(Figure 1B)**. The results showed that high cell density resulted in Jurkat T cell activation, while low cell density did not. This indicated that the secretion of ADA1.CD3scFv can be triggered by cellular stress, while ADA1.CD3scFv is not secreted in physiological conditions. To localize the ADA1.CD3scFv protein, ADA enzyme activity assays were also performed on the Jurkat T cell culture media or cells **(Figure 1C)**. The result showed that the activity of the ADA in the cell culture medium was not increased by high cell density. However, the activity of the ADA on the surface of Jurkat T cell was significantly increased by high cell density, but not by low cell density. This indicated that ADA1.CD3scFv was located on the surface of the transfected Jurkat T cells.

We then investigated whether ADA1.CD3scFv activates human CAR T cells only in stress conditions and not under physiological conditions. To generate MR-CAR T cells, human PBMCs (peripheral blood mononuclear cells) were co-transduced with two RD114-pseudotyped retroviral vectors. The first MR vector encoded CD26 and ADA1.CD3scFv, while the second CAR vector encoded either a GPC3- or HER2-specific CAR **(sFigure 1A)**. Cell surface expression on the MR-CAR T Cells of the CARs and CD26 was detected by flow cytometry **(sFigure 1B-D)**. After transduction, both HER2 and GPC3 CARs were stably expressed on the surface of human peripheral blood T cells. The expression level of CD26 on both HER2- or GPC3 MR-CAR T cells increased significantly compared to that on HER2- or GPC3-specific CAR T cells, respectively. The transcriptional expression of ADA1.CD3scFv was demonstrated by western blotting **(sFigure 1E).**

Next, we investigated whether human CAR T cells are exclusively activated by ADA1.CD3scFv under stress conditions and not in physiological ones. We used qRT-PCR to assess the transcriptional expression of interferon-gamma (IFNγ) in HER2-specific or GPC3-specific MR-CAR T cells cultured under physiological conditions. In these cell ADA1.CD3scFv did not stimulate the expression of IFN-γ automatically **(Figure 1D)**. Furthermore, RNA sequencing showed that MR-CAR T cells did not exhibit an enhanced inflammatory gene prolife **(Figure 1E).**

The next step was to determine whether MR-CAR T cells were activated upon engaging tumor cells. The HER2-CAR T cells were cultured in the presence or absence of HER2-positive NSCLC A549 tumor cells at a 1:1 ratio for 24 hours. The culture media were analyzed by an IFN-γ ELISA. In the absence of tumor cells, ADA1.CD3scFv in HER2-CAR T cells did not induce IFN-γ expression, but the presence of the HER2-positive NSCLC 459 cell induced significantly increased the amounts of IFN-γ production by the HER2-MR-CAR T cell **(Figure 1F)**. The results are consistent with HER2-MR-CAR T cells showing increased expression of T-bet, granzyme B, IFN-γ, and IL-2 upon CD3 antibody stimulation (**sFigure 2A-B**). These findings suggested that engagement with tumor cells could trigger the release of ADA1.CD3scFv, which subsequently acts as a trans-signal to activate CAR T cells in a tumor antigen-specific manner.

ADA1 activates CD26, which is expressed on various cell types, including T cells^30,36,38,47^. Consequently, we wondered if a membrane-bound version of ADA1 could result in autocrine activation of CAR T cells through interacting with CD26. HER2-CAR-T cells or GPC3-CAR-T cells co-expressing membrane-bound ADA1 and CD26 were cultured with or without HER2-positive A549 NSCLC or GPC3-positive GPC3 HCC overnight. An ELISA assay revealed that membrane-bound ADA1 induced the expression of IFN-γ in both HER2-CAR-T cells or GPC3-CAR-T cells independent of tumor stimulation, with tumor stimulation further increasing IFN-γ expression **(Figure 1G)**. This indicated that membrane-bound ADA1 induces autocrine activation of human T cells independent of tumor antigens. Taken together, these findings demonstrate that the ADA1.CD3scFv construct optimize ADA1 in T cell-based therapies.

### ADA1.CD3scFv enhances CAR T cell expansion preferentially without impacting tumor cells

CD26 on T cells functions as an extracellular ADA1 receptor, and its expression is elevated following T cell activation. However, ADA1 has a relatively low affinity for CD26 (K_D_=1.8^−8^M)^48^, thus we conducted assessments to determine the efficacy of ADA1-CD26 binding, specifically by measuring the presence of ADA1 on the surface of CD26-positive Jurkat T cells. The results indicated that the addition of extra ADA1 did not result in increased binding to the CD26-positive Jurkat T cells, suggesting an ineffective binding of ADA1 to CD26 (**sFigure 3A**).

We then created a CD26-positive NFAT or NF-_K_B luciferase reporter human Jurkat T cell line using a retroviral vector generating human CD26 in order to ascertain if the CD3scFv is required for anchoring ADA1 to CAR T cells. After being transduced with the retroviral vector expressing ADA1.CD3scFv or ADA1, human Jurkat T cells with NFAT or NF-_K_B luciferase reporter were subsequently grown at high cell density to promote the secretion of ADA1.CD3scFv or ADA1. The amount of ADA1 attached to the Jurkat T cell membrane was assessed using an ADA enzyme activity assay after the Jurkat T cells had been in culture for 24 hours. According to the findings, ADA1 only had approximately 40% of the activity on CD26-positive Jurkat T cells compared to ADA1.CD3scFv (**Figure 2A-B**). However, the ADA1.CD3scFv’s activity on Jurkat T cells that lacked CD26 was comparable to that of the ADA1.CD3scFv on CD26-positive Jurkat T cells. The results aligned with flow analysis of ADA1 on the surface of Jurkat T cells **(sFigure 3B)** and the ADA activity assay conducted on HER2-specific MR-CAR T cells **(sFigure 3C**). These findings demonstrated the critical role of the CD3scFv in strengthening the interaction between ADA1.CD3scFv and human T cells.

**Figure 2.**
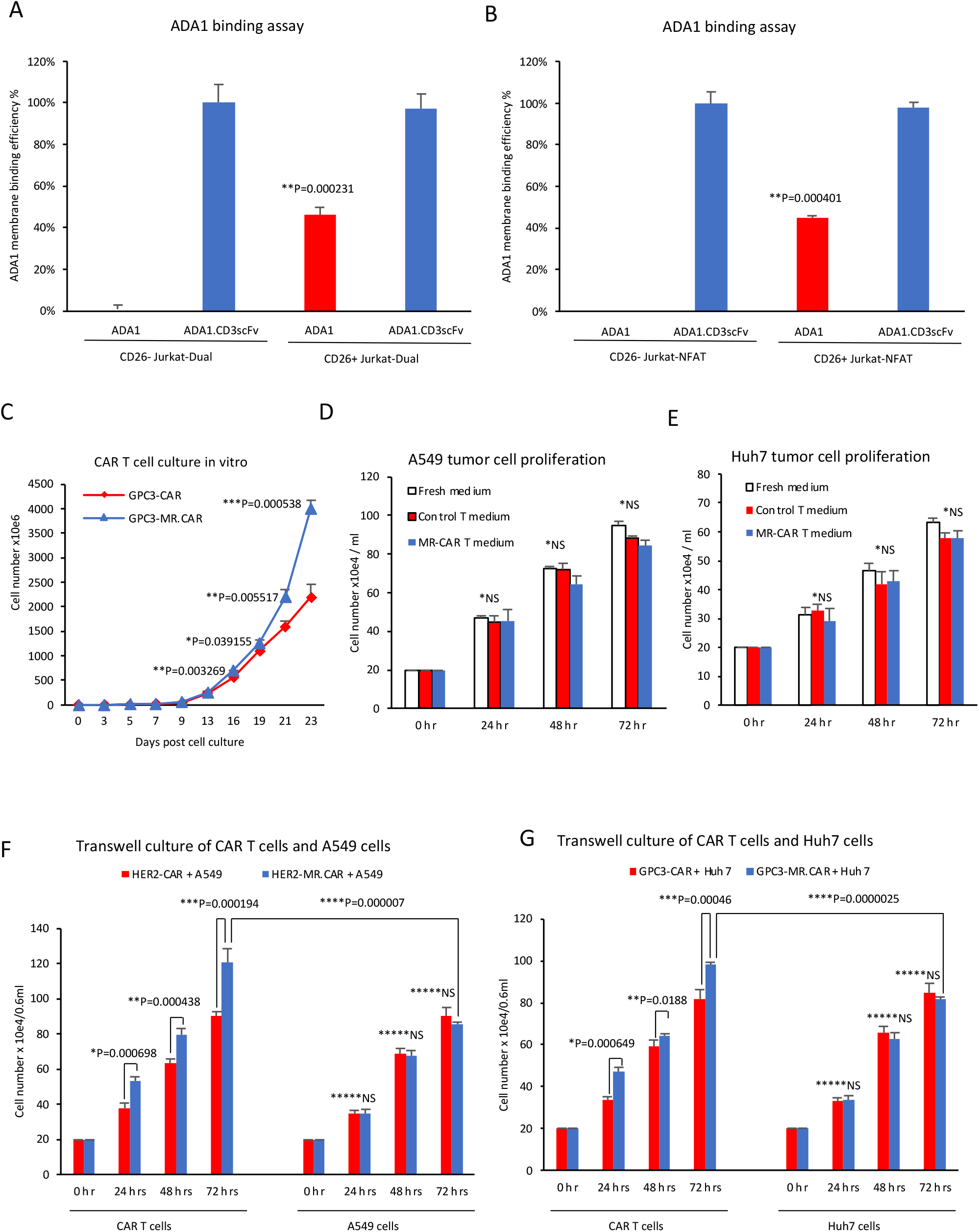
ADA1.CD3scFv enhances CAR T cell expansion preferentially without impacting tumor cells. **(A-B)** Luciferase reporter Jurkat-Dual or Jurkat-NFAT cells either expressing CD26 or lacking CD26 were transduced with overexpressing vectors of ADA1 or ADA1.CD3scFv and cultured for 24 hours. After incubation, the cells were collected and subjected to an ADA activity assay to determine the amount of ADA1 on the surface of the cells. CD26-positive Jurkat T cells transduced with ADA1.CD3scFv overexpressing vector were used as maximum value to calculate the percentage. The data were presented as the percentage of ADA1 or ADA1.CD3scFv binding with the Jurkat T cells. The figure indicated the P values for the binding of ADA1 with CD26+ Jurkat T cells verse the binding of ADA1.CD3scFv with CD26+ Jurkat T cells. **(C)** GPC3-MR-CAR T cells and GPC3-CAR T cells were expanded in vitro, and the cell numbers were determined at different time points using a hemocytometer. The results were presented as a growth curve. The figure indicated the P values for GPC3-MR-CAR vs GPC3-CAR. **(D-E)** The conditioned medium form HER2-MR-CAR T cells or HER2-CAR T cells was added to A549 cell cultures and incubated for 72 hours. Similarly, the conditioned medium form GPC3-MR-CAR T cells or GPC3-CAR T cells was added to Huh7 cell culture and incubated for 72 hours. Fresh medium was used as control. The numbers of tumor cell were quantified daily using a hemocytometer. *NS for HER2-MR-CAR vs HER2-CAR, and GPC3-MR-CAR vs GPC3-CAR. **(F-G)** HER2-MR-CAR T cells or HER2-CAR T cells were co-cultured with A549 tumor cells in a transwell plate for 72 hours. Similarly, GPC3-MR-CAR T cells or GPC3-CAR T cells were co-cultured with Huh7 tumor cells in a transwell plate for 72 hours. The CAR T cells and tumor cells were counted using a hemocytometer and presented. The figure indicated the P values. The data are presented as mean ± SD, n = 3.

The impact of ADA1.CD3scFv on T cell survival and proliferation was the subject of our next investigation. Our findings demonstrated that at day 23 of in vitro growth, the number of GPC3-specific MR-CAR T cells was two times higher than that of conventional GPC3-CAR T cells (**Figure 2C**). Given the possibility that tumor cells could utilize inosine as an energy source, we also looked at whether ADA1.CD3scFv production by CAR T cells could be advantageous to tumor cells. However, we found that the culture medium from MR-CAR T cells did not promote tumor growth **(Figure 2D-E)**. To further examine the effect on tumor cells we conducted a transwell cell culture experiment using HER2-MR-CAR T cells or HER2-CAR T cells with A549, as well as GPC3-MR-CAR T cells or GPC3-CAR T cells with Huh7 cells (**Figure 2F-G**). The MR-CAR T cells proliferated more quickly than CAR T cells in both tests. Tumor cell proliferation did not differ between co-culture settings. This indicated that ADA1.CD3scFv selectively increases CAR T cell growth without affecting tumor cells. This is most likely because the ADA1 is mostly found in the cytoplasm, and once secreted, it binds to CAR T cells again and converts adenosine to inosine in the immediate vicinity of the CAR T cells.

### Overexpression of CD26 resisted TGF-β1 suppression and promoted CAR T cell mobility

The expression of CD26 is carefully regulated during T lymphocyte development. TGF-β1 is a component of the regulatory system and inhibits CD26 expression^39,40^. We therefore examined how constitutive CD26 expression induced by the MR helper vector might change CD26 expression when TGF-β1 was present. Using flow cytometry, CD26 cell surface expression was evaluated. Human T cells that had undergone transduction expressed CD26 at a level that was noticeably greater than non-transduced (NT) human T cells (**Figure 3A**). After 48 hours of in vitro incubation in the presence of 20ng/ml of TGF-β1, CD26 expression on NT T cells was downregulated by 40%, whereas MR vector-transduced T cells defied TGF-β1-mediated suppression of CD26 expression.

**Figure 3.**
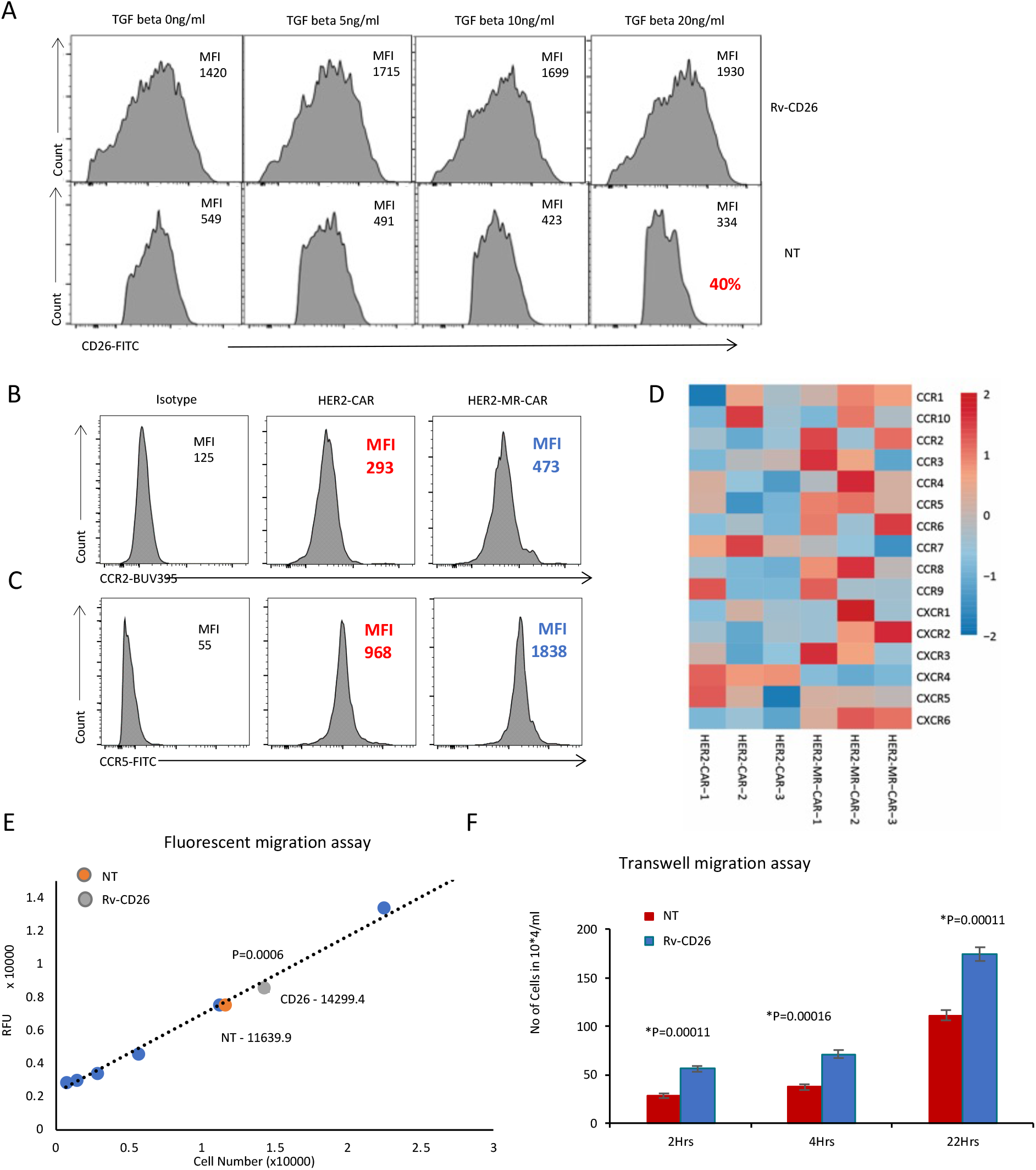
Overexpression of CD26 resisted TGFβ1 suppression and promoted CAR T cell mobility. **(A)** Human PBMC were activated using anti-CD3/CD28 antibodies and transduced with CD26-overexpressing vector and expanded in vitro. Non-transduced T cells (NT) were used as a negative control. The T cells were then cultured in the presence of TGFβ1 at the indicated concentrations for 48 hours, and CD26 expression was detected by flow cytometry. **(B-C)** HER2-MR-CAR T cells or HER2-CAR T cells were stained with antibodies against human CCR2 and CCR5, followed by flow cytometric analysis. **(D)** The heatmap shows the expression levels of chemokine receptor genes in two groups, represented on the Y-axis. The expression levels were converted to Z-scores to enable comparison between the different chemokine receptor genes. Triplicate samples (N=3) were used for each group and are represented on the X-axis. The heatmap provides a visual representation of the relative expression levels of chemokine receptor genes in the two groups, with warmer colors indicating higher expression levels and cooler colors indicating lower expression levels. **(E-F)** Human T cells transduced with CD26 overexpressing vector and control T cells were subjected to a fluorescent migration assay and transwell migration assay to determine their migration capacity. The figure indicated the P values for human T cells transduced with CD26 overexpressing vector verse NT.

We questioned if MR-CAR T cells would express increased amounts of chemokine receptors given that human CD26^high^ T cells strongly induce immune responses against numerous forms of malignancy through enhanced migration^38^. Using flow cytometry, the expression of CCR2 and CCR5 on T cell surfaces was examined. When compared to HER2-CAR T cells, we found that HER2-MR-CAR T cells have considerably higher levels of CCR2 and CCR5 expression (**Figure 3B-C**). This demonstrated that on HER2-CAR T cells, the expression of CD26 increases the production of CCR2 and CCR5. Additionally, mRNA sequencing analysis proved that HER2-MR-CAR T cells have greater chemokine receptor expression profiles (**Figure 3D**). Using a fluorescent migration test and a transwell migration assay, the mobility of these T cells was examined (**Figure 3E–F**). At 2, 4, and 22 hours after transduction with the MR retroviral vector, T cells showed improved migration. In conclusion, CD26 expression increases T cell mobility and resists TGF-β1 inhibition, demonstrating benefits of overexpressing CD26 in CAR T cells.

### MR-CAR T cells displayed enhanced antitumor cytotoxicity in vitro

To compare the effectiveness of HER2-MR-CAR T cell and HER2-CAR T cells to kill tumor cells, we quantified the release of lactate dehydrogenase (LDH) in the media at various effector to target ratios. As a negative control, non-transduced T cells (NT) were used as effectors. At ratios of 1:1, 5:1, 10:1, or 20:1, HER2-MR-CAR T cells or HER2-CAR T cells were co-cultured with HER2^high^ Calu3 (NSCLC) **(Figure 4A-C)** and HER2^low^ A549 (NSCLC), respectively **(Figure 4D-F)**. After 4 hours of culture at a ratio of 10:1 or above and after 18 hours of culture at a ratio of 1:1 or above, the HER2-CAR T cells eliminated the HER2-positive Calu3 and A549, demonstrating the potency of the HER2-specific CAR. We found no difference in the capacity of HER2-CAR T cells to kill Calu3 tumor cells that were HER2^high^ compared to A549 tumor cells that were HER2^low^. This shows that both HER2^high^ and HER2^low^ tumors can be killed by HER2-CAR T cells. The HER2-MR-CAR T cells demonstrated improved cytotoxicity against both Calu3 and A549 after 4 hours of culture at a ratio of 10:1 or higher and after 18 hours of culture at a ratio of 5:1 or higher. This suggests that the expression of CD26 and ADA1.CD3scFv enhances the ability of HER2-CAR T cells to kill targeted tumor cells.

**Figure 4.**
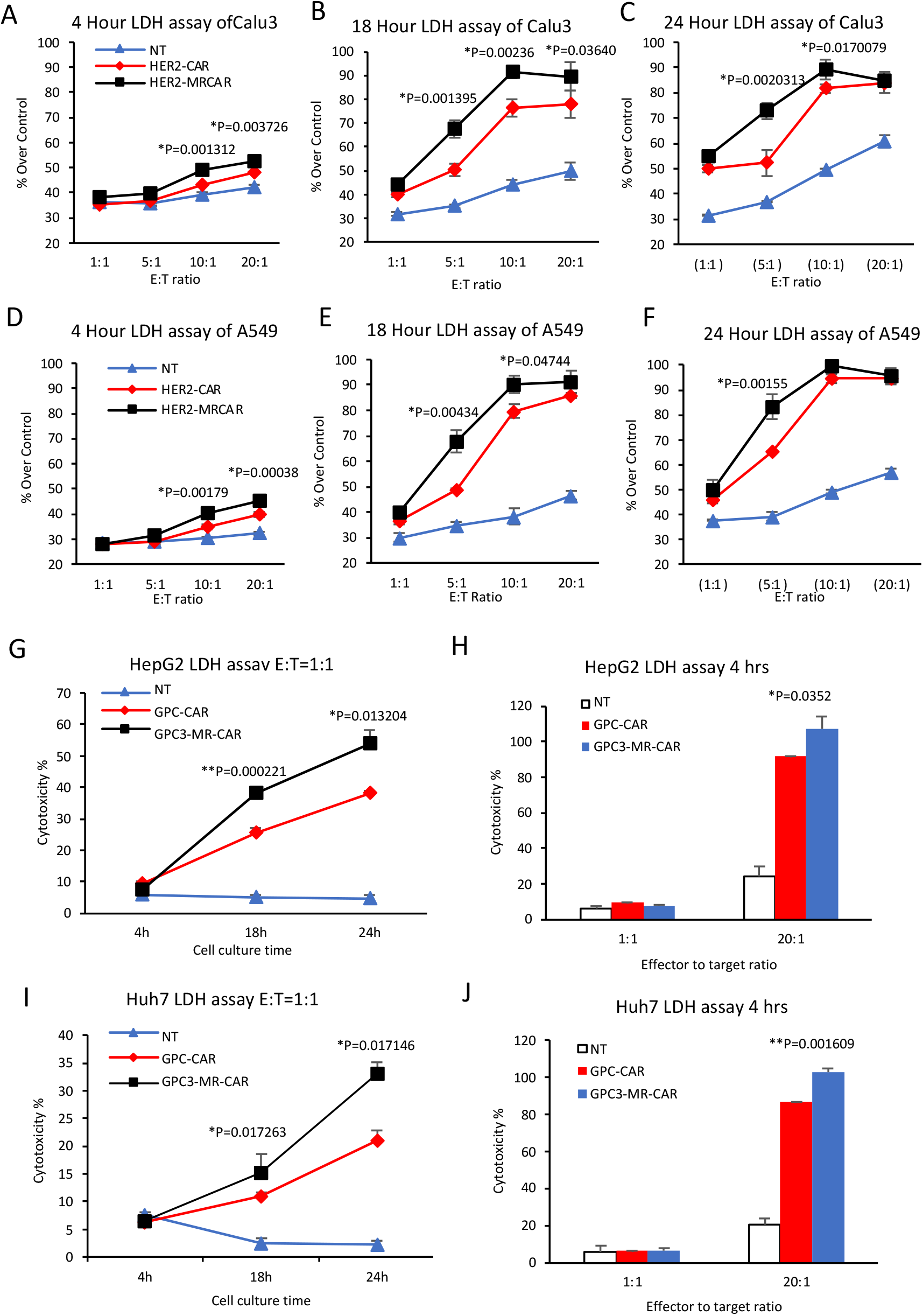
MR-CAR T cells displayed enhanced antitumor cytotoxicity in vitro. **(A-F)** Human PBMC were activated by anti-CD3/CD28 antibodies and transduced with either HER2-MR-CAR or HER2-CAR and expanded in vitro. Non-transduced T cells (NT) were used as a negative control. The cytotoxic activity of HER2-MR-CAR and HER2-CAR T cells against Calu3 and A549 cells was evaluated using a lactate dehydrogenase (LDH) assay. The percentage of cytotoxicity was calculated as the ratio of LDH release in the effector-target cell co-culture relative to the maximum LDH release control. The figure indicated the P values for HER2-MR-CAR verse HER2-CAR. **(G-J)** Human PBMC were activated by anti-CD3/CD28 antibodies and transduced with either GPC3-MR-CAR or GPC3-CAR and expanded in vitro. Non-transduced T cells (NT) were used as a negative control. The cytotoxic activity of GPC3-MR-CAR and GPC3-CAR T cells against HepG2 and Huh7 cells was evaluated using a lactate dehydrogenase (LDH) assay. The figure indicated the P values for GPC3-MR-CAR verse GPC3-CAR.

We next investigated GPC3-MR-CAR T cell’s cytotoxicity against GPC3-positive HCC Huh7 and HepG2. GPC3-MR-CAR T cells or GPC3-CAR T cells were co-cultured with Huh7 **(Figure 4G-H)** or HepG2 **(Figure 4I-J)** at ratios of 1:1 or 20:1. The GPC3-CAR T cells killed both Huh7 and HepG2, which were GPC3-positive, after 4 hours of culture at a ratio of 20:1 and after 18 hours of culture at a ratio of 1:1, indicating the effectiveness of the GPC3-specific CAR. The GPC3-MR-CAR T cells displayed enhanced cytotoxicity against both Huh7 and HepG2 after 4 hours of culture at a ratio of 20:1 and after 18 hours of culture at a ratio of 1:1. To explore the effects of PD-L1 inhibition on GPC3-MR-CAR T cell proliferation and effector functions, we assessed the expression of PD-1 on the surface of both GPC3-MR-CAR T cells and GPC3-CAR T cells (**sFigure 4A**). Interestingly, both cell types exhibited increased PD-1 expression. However, while PD-L1 significantly suppressed the proliferation of GPC3-CAR T cells, GPC3-MR-CAR T cells showed significant rescue in their proliferation (**sFigure 4B**). Additionally, we found that PD-L1 did not inhibit the expression of T-bet, granzyme B, IL-2, IFN-γ, and TNF-α in GPC3-CAR T cells, whereas GPC3-MR-CAR T cells exhibited enhanced expression levels of these effector molecules (**sFigure 4C-J**). These results suggest that the expression of CD26 and ADA1.CD3scFv enhances the ability of GPC3-CAR T cells to kill targeted tumor cells and confers resistance to PD-L1 inhibition.

We next asked whether the enhanced cytotoxicity of HER2-MR-CAR T cells could result in T cell exhaustion. To this end, we conducted mRNA profiling and found that HER2-specific MR-CAR T cells exhibited a lower level of T cell exhaustion markers compared to HER2-CAR T cells **(sFigure 5**). Furthermore, mRNA gene expression analysis revealed that HER2-specific MR-CAR T cells displayed increased NOTCH signaling, enhanced DNA repair mechanisms, and complement activation compared to HER2-CAR T cells (**sFigure 6A-B**, **table 1**). Notch signaling is known to regulate cytokine and chemokine expression, cell migration, and promotes cell survival through anti-apoptotic protein expression (Bcl-2, Bcl-xL, Mcl-1), and activation of the PI3K/Akt and MAPK signaling pathways involved in cell survival and proliferation^49–53^. The intracellular complement system is involved in metabolic reprogramming necessary for effector responses^54^. In addition, HER2-MR-CAR T cells have a reduced inflammatory response (i.e., TGF-β1 & IFN-α) compared to HER2-CAR T cells, as well as reduced levels of pro-inflammatory cytokines such as IL6, IFN-g, TNF-a, and IL2. This may be beneficial in preventing unwanted immune responses and promoting a more controlled immune response against tumor cells. HER2-CAR T cells showed an increased expression of genes related to glycolysis, ROS, oxidative phosphorylation, UV-response, hypoxia, and apoptosis. This suggests that these cells may experience cellular stress and could be at risk of damage or death due to the potentially harmful effects of these processes. HER2-CAR T cells also showed an upregulation of genes related to MTORC1, P53, and KRAS, which may suggest a potential for T cell exhaustion. In conclusion, HER2-MR-CAR T cells exhibit reduced exhaustion and display advanced cell division phenotypes.

**Table 1.**
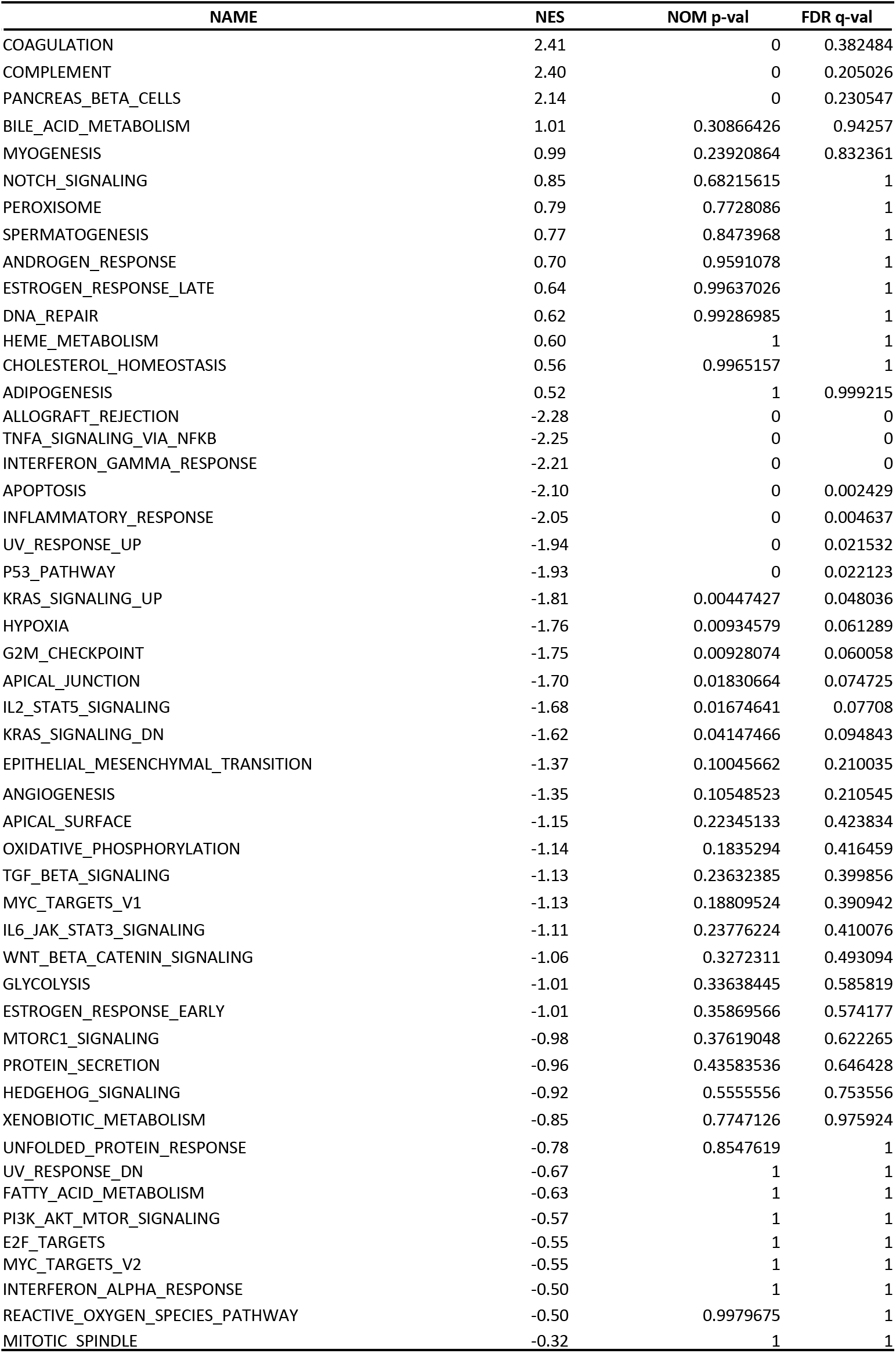
mRNA sequencing analysis NES, p value and FDR q-val. The table summarizes the normalized enrichment scores (NES), p-values, and false discovery rate q-values (FDR q-val) for genes and pathways that were differentially expressed between HER2-MR-CAR T cells and HER2-CAR T cells based on mRNA sequencing analysis. Higher NES values represent greater differential expression between the two groups.

| NAME | NES | NOM p-val | FDR q-val |
| --- | --- | --- | --- |
| COAGULATION | 2.41 | 0 | 0.382484 |
| COMPLEMENT | 2.40 | 0 | 0.205026 |
| PANCREAS_BETA_CELLS | 2.14 | 0 | 0.230547 |
| BILE_ACID_METABOLISM | 1.01 | 0.30866426 | 0.94257 |
| MYOGENESIS | 0.99 | 0.23920864 | 0.832361 |
| NOTCH_SIGNALING | 0.85 | 0.68215615 | 1 |
| PEROXISOME | 0.79 | 0.7728086 | 1 |
| SPERMATOGENESIS | 0.77 | 0.8473968 | 1 |
| ANDROGEN_RESPONSE | 0.70 | 0.9591078 | 1 |
| ESTROGEN_RESPONSE_LATE | 0.64 | 0.99637026 | 1 |
| DNA_REPAIR | 0.62 | 0.99286985 | 1 |
| HEME_METABOLISM | 0.60 | 1 | 1 |
| CHOLESTEROL_HOMEOSTASIS | 0.56 | 0.9965157 | 1 |
| ADIPOGENESIS | 0.52 | 1 | 0.999215 |
| ALLOGRAFT_REJECTION | -2.28 | 0 | 0 |
| TNFA_SIGNALING_VIA_NFKB | -2.25 | 0 | 0 |
| INTERFERON_GAMMA_RESPONSE | -2.21 | 0 | 0 |
| APOPTOSIS | -2.10 | 0 | 0.002429 |
| INFLAMMATORY_RESPONSE | -2.05 | 0 | 0.004637 |
| UV_RESPONSE_UP | -1.94 | 0 | 0.021532 |
| P53_PATHWAY | -1.93 | 0 | 0.022123 |
| KRAS_SIGNALING_UP | -1.81 | 0.00447427 | 0.048036 |
| HYPOXIA | -1.76 | 0.00934579 | 0.061289 |
| G2M_CHECKPOINT | -1.75 | 0.00928074 | 0.060058 |
| APICAL_JUNCTION | -1.70 | 0.01830664 | 0.074725 |
| IL2_STAT5_SIGNALING | -1.68 | 0.01674641 | 0.07708 |
| KRAS_SIGNALING_DN | -1.62 | 0.04147466 | 0.094843 |
| EPITHELIAL_MESENCHYMAL_TRANSITION | -1.37 | 0.10045662 | 0.210035 |
| ANGIOGENESIS | -1.35 | 0.10548523 | 0.210545 |
| APICAL_SURFACE | -1.15 | 0.22345133 | 0.423834 |
| OXIDATIVE_PHOSPHORYLATION | -1.14 | 0.1835294 | 0.416459 |
| TGF_BETA_SIGNALING | -1.13 | 0.23632385 | 0.399856 |
| MYC_TARGETS_V1 | -1.13 | 0.18809524 | 0.390942 |
| IL6_JAK_STAT3_SIGNALING | -1.11 | 0.23776224 | 0.410076 |
| WNT_BETA_CATENIN_SIGNALING | -1.06 | 0.3272311 | 0.493094 |
| GLYCOLYSIS | -1.01 | 0.33638445 | 0.585819 |
| ESTROGEN_RESPONSE_EARLY | -1.01 | 0.35869566 | 0.574177 |
| MTORC1_SIGNALING | -0.98 | 0.37619048 | 0.622265 |
| PROTEIN_SECRETION | -0.96 | 0.43583536 | 0.646428 |
| HEDGEHOG_SIGNALING | -0.92 | 0.5555556 | 0.753556 |
| XENOBIOTIC_METABOLISM | -0.85 | 0.7747126 | 0.975924 |
| UNFOLDED_PROTEIN_RESPONSE | -0.78 | 0.8547619 | 1 |
| UV_RESPONSE_DN | -0.67 | 1 | 1 |
| FATTY_ACID_METABOLISM | -0.63 | 1 | 1 |
| PI3K_AKT_MTOR_SIGNALING | -0.57 | 1 | 1 |
| E2F_TARGETS | -0.55 | 1 | 1 |
| MYC_TARGETS_V2 | -0.55 | 1 | 1 |
| INTERFERON_ALPHA_RESPONSE | -0.50 | 1 | 1 |
| REACTIVE_OXYGEN_SPECIES_PATHWAY | -0.50 | 0.9979675 | 1 |
| MITOTIC_SPINDLE | -0.32 | 1 | 1 |

### MR-CAR T cells have enhanced antitumor activities in multiple xerograph mouse models

We hypothesized that HER2-specific or GPC3-specific MR-CAR T cells may inhibit HER2-positive or GPC3-positive tumor growth more effectively than control CAR T cells in xenograft animal models based on the above-described in vitro results. Using a subcutaneous A549 tumor model, we first looked into the anticancer effects of HER2-specific CAR T cells. 2×10^6^ A549 cells were injected subcutaneously (s.c.) into the right flank of NSG mice to create the tumors. The mice received 2×10^6^ HER2-MR-CAR T cells, 2×10^6^ HER2-CAR T cells, or phosphate-buffered saline (PBS), respectively, once the tumor size reached an average of 4-6 mm in diameter. Comparative to control animals, HER2-CAR T cells slightly reduced the development of the A549 tumor. In contrast, mice that received HER2-MR-CAR T cells demonstrated a considerably greater inhibition of A549 tumor growth compared to mice that received HER2-CAR T cells or PBS (**Figure 5A**). Additionally, the mice’s body weight was examined, and neither the administration of HER2-MR-CAR T cells nor the administration of HER2-CAR T cells was associated with any changes in body weight (**Figure 5B**). The effectiveness of HER2-MR-CAR T cells in treating HER2^high^ human NSCLC Calu3 xenograft mice model was next assessed. Tumors treated with HER2-MR-CAR T cells showed a trend toward reduced size when compared to unmodified HER2-CAR T cells after a single dosage of 2×10^6^ CAR T cells (**sFigure 7**). These findings indicate that HER2-MR-CAR T cells boost antitumor effectiveness without causing any obvious adverse effects.

**Figure 5.**
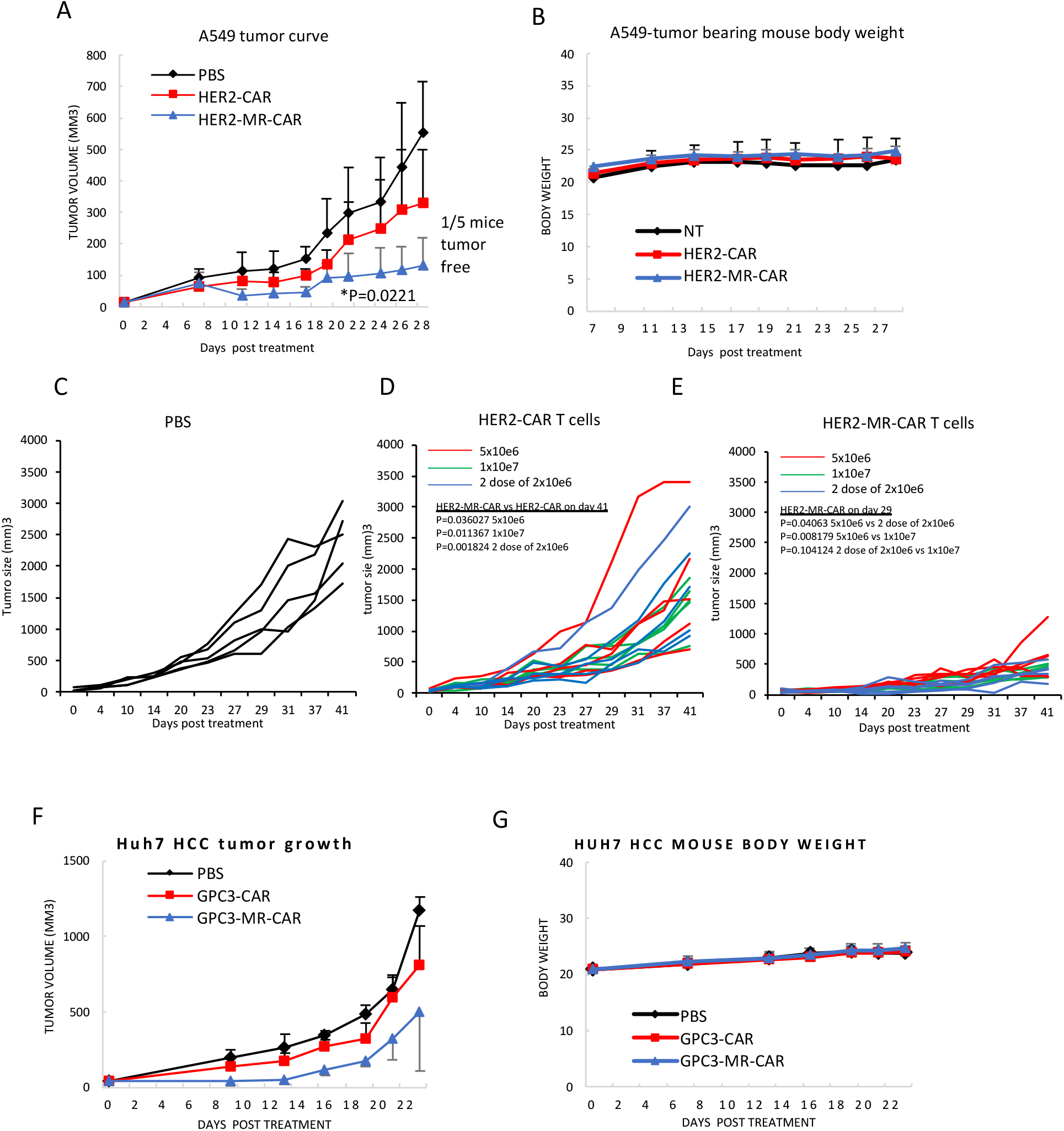
MR-CAR T cells demonstrated anti-tumor activity in xenograft mouse models. **(A-B)** A murine melanoma xenograft model was established in NSG mice by subcutaneous inoculation of 2 × 10^6^ A549 tumor cells on the right flank. When the average tumor size reached 4-6 mm in diameter, experimental mice were treated with a single dose of either 2 × 10^6^ HER2-MR-CAR T cells, 2 × 10^6^ HER2-CAR T cells, or PBS. Tumor size and mouse body weight were monitored every two to three days. Data represent mean ± s.d. (n = 5). The results showed that treatment with HER2-MR-CAR T cells inhibited A549 tumor growth more effectively compared to treatment with HER2-CAR T cells (P =0.0327, P=0.0258, and P=0.0221 for HER2-MR-CAR verse HER2-CAR on day 20, day 22, and day 24 respectively). No difference in body weight was observed among groups. **(C-E)** A549 tumor bearing mice were treated with a single dose of either 5 × 10^6^ or 1 × 10^7^ or two doses of 2 × 10^6^ (at one-week intervals) HER2-MR-CAR T cells, a single dose of either 5 × 10^6^ or 1 × 10^7^ or two doses of 2 × 10^6^ (at one-week interval) HER2-CAR T cells, or PBS. Tumor size and mouse body weight were monitored every two to three days. Data represent mean ± s.d. (n = 5). The figure indicates the P values. No difference in body weight was observed among groups. **(F-G)** A murine melanoma xenograft model was established in NSG mice by subcutaneous inoculation of 2 × 10^6^ Huh7 tumor cells on the right flank. When the average tumor size reached 4-6 mm in diameter, experimental mice were treated with a single dose of either 2 × 10^6^ GPC3-MR-CAR T cells, 2 × 10^6^ GPC3-CAR T cells, or PBS. Tumor size and mouse body weight were monitored every two to three days. Data represent mean ± s.d. (n = 5). The results showed that treatment with GPC3-MR-CAR T cells resulted in a trend towards reduced tumor growth compared to treatment with GPC3-CAR T cells (P=0.13 and P=0.14 for GPC3-MR-CAR T cells vs GPC3-CAR T cells on day 13 and day 16 respectively). No difference in body weight was observed among groups.

Then, by increasing the dose, we investigated the anticancer effects of HER2-MR-CAR T cells. 5×10^6^HER2-MR-CAR T cells, 10×10^6^ HER2-MR-CAR T cells, or two doses of 2×10^6^ HER2-MR-CAR T cells were given to groups of mice in subcutaneous A549 animal models. As a control, HER2-CAR-T cells or PBS were used. The findings demonstrated that HER2-MR-CAR T cells were superior to HER2-CAR T cells at inhibiting tumor growth at the same dose (**Figure 5C-E**). Additionally, compared to 5×10^6^ HER2-MR-CAR T cells, 10×10^6^ HER2-MR-CAR T cells or two doses of 2 × 10^6^ HER2-MR-CAR T cells showed an improved reduction of tumor growth. In mice receiving either therapy, there was no indication that their body weight had changed. These findings imply that HER2-MR-CAR T cells’ anticancer activities can be further enhanced by increasing the dose or undergoing more treatments without manifesting any obvious negative side effects.

We also sought to replicate the improved antitumor effects of MR-CAR T cell with GPC3-specific CAR in a mouse model of GPC3-positive hepatocellular carcinoma (HCC). To establish the tumors, 2 × 10^6^ Huh7 cells were inoculated subcutaneously into the right flank of NSG mice. Once the tumor size reached an average of 4-6 mm in diameter, the mice were treated with either 2 × 10^6^ GPC3-MR-CAR T cells, 2 × 10^6^ GPC3-CAR T cells, or PBS delivered via the tail veil. In comparison to mice treated with GPC3-CAR T cells or PBS, tumor progression in the GPC3-MR-CAR-T cell-treated mice were considerably reduced **(Figure 5F)**. The mice receiving either form of CAR T cell therapy showed no indication that their body weight had changed (**Figure 5G**). These findings imply that GPC3-specific MR-CAR T cells improve antitumor effectiveness without causing any significant side effects. In conclusion, our research indicates that in preclinical mouse models, MR-CAR increases the effectiveness of CAR T cell therapy.

### MR-CAR T cells retained their capability to proliferate, migrate, and lyse tumor cells in the tumor microenvironment (TME)

Subsequently, we examined the T cell responses within the tumor microenvironment. A549 tumor-bearing mice were received 10×10^6^ of either HER2-MR-CAR T cells, HER2-CAR T cells, or non-transduced T cells. At specific times after T cell administration, tumor tissues were dissected, and single-cell suspensions were prepared. The concentration of inosine in the single-cell washout was determined using an inosine assay **(Figure 6A)**. The results show that the concentration of inosine significantly increased in the HER2-MR-CAR group compared to that in HER2-CAR or non-transduced group, while there was no difference in inosine concentration between the HER2-CAR and non-transduced group. This indicated that ADA1.CD3scFv was expressed in the tumor microenvironment and effectively converted adenosine to inosine in the tumor microenvironment.

**Figure 6.**
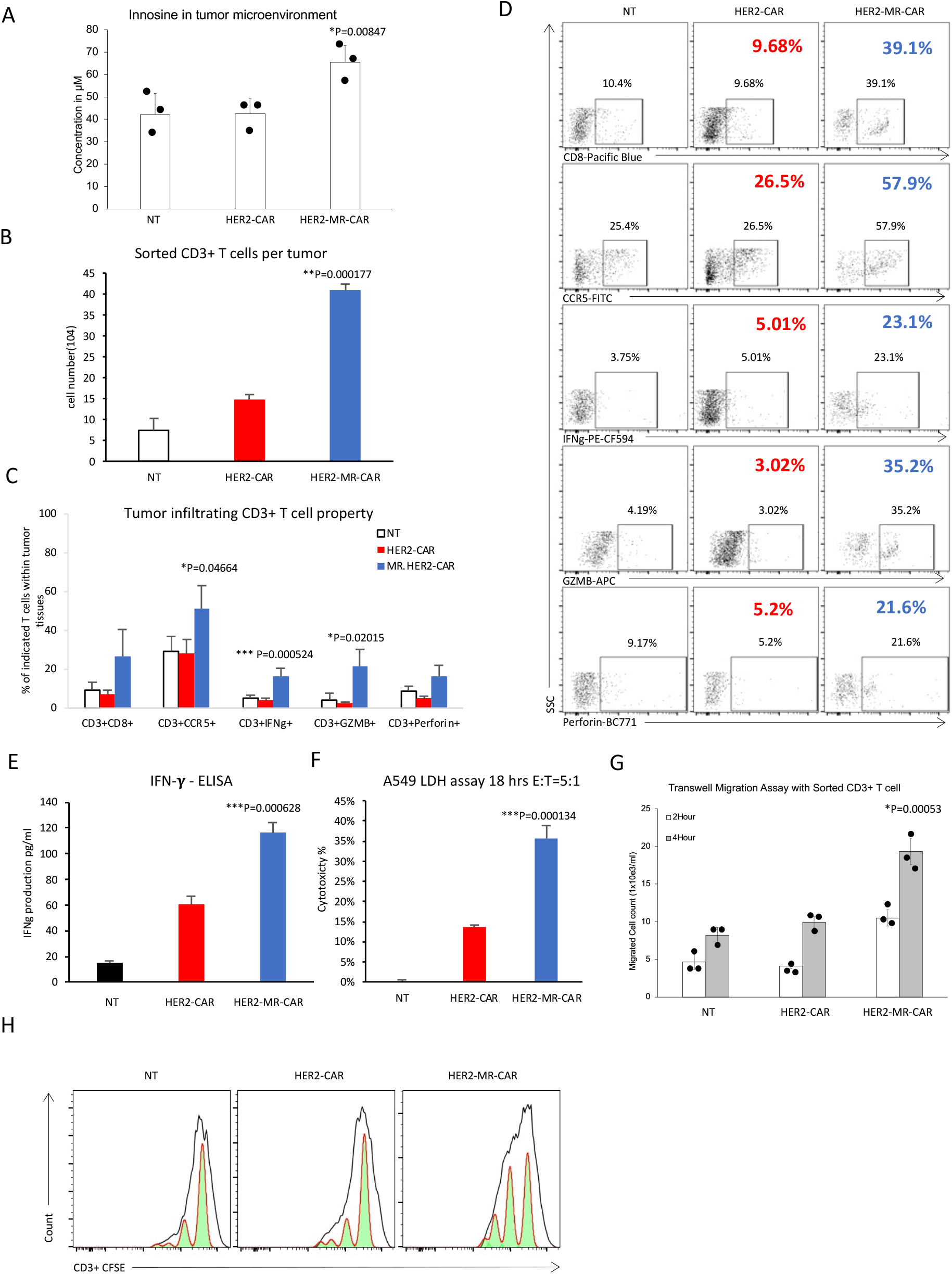
MR-CAR T cells retained their capability to proliferate, migrate, and lyse tumor cells in the tumor microenvironment (TME). A murine melanoma xenograft model was established in NSG mice by subcutaneous inoculation of 2 × 10^6^ A549 tumor cells on the right flank. When the average tumor size reached 200-300 mm^3^, experimental mice were treated with a single dose of either 1 × 10^7^ HER2-MR-CAR T cells, 1 × 10^7^ HER2-CAR T cells, or Non-transduced T cells (NT). **(A)** Seven days after treatment, tumor tissues were dissected, and a single-cell suspension was prepared, with the cell wash buffer then subjected to an ADA activity assay to measure inosine concentrations. *P=0.00847 for HER2-MR-CAR verse HER2-CAR. **(B)** After seven days of treatment, single cell suspensions were prepared from each tumor tissue, and CD3+ cells were sorted using flow cytometry. The number of sorted cells was quantified and presented (n=3). *P=0.000177 for HER2-MR-CAR verse HER2-CAR. **(C-D)** Single cell suspensions from tumor tissues were stained with antibodies against CD3, CD8, CCR5, IFN-γ, Granzyme B, and Perforin and analyzed by flow cytometry. The resulting data were presented as a flow cytometry dot plot, and the gated cell populations were quantified and presented in a column figure. The figure indicated P value. **(E-F)** Sorted CD3+ cells were co-cultured with A549 tumor cells at an E:T ratio of 1:1 overnight. After incubation, the culture medium was collected and subjected to an ELISA to measure the expression of IFN-γ. To determine the tumor-killing capacity, a LDH assay was performed. The figure indicated P value. **(G)** Sorted CD3+ cells were co-cultured with A549 tumor cells in a transwell culture plate to assess the migration capacity of the CD3+ cells. *P=0.00053 for HER2-MR-CAR verse HER2-CAR. **(H)** Sorted CD3+ cells were labeled with CFSE and then co-cultured with A549 tumor cells at an E:T ratio of 1:1 for 48 hours. After incubation, the cells were collected, stained with anti-CD3 antibody, and analyzed by flow analysis to determine their proliferation.

Mice treated with the HER2-MR-CAR T cells had a higher number of tumor-infiltrating T cells compared to those receiving HER2-CAR T cells or non-transduced T cells **(Figure 6B)**. A higher number of CD8+ TILs was observed in the tumor tissue of mice receiving HER2-MR-CAR T cells (**Figure 6C-D)**. Furthermore, the treatment with HER2-MR-CAR T cells resulted in an increased number of CD3+ TILs expressing CCR5, IFNγ, granzyme B, and perforin compared with HER2-CAR T cell treatment. In contrast, HER2-CAR T cells showed no difference compared to non-transduced T cells, suggesting a significant immunosuppression of their immunological activity in the tumor microenvironment. As indicated by the increased number of tumor-infiltrating T cells, and the expression of key activation markers, these results suggest that the HER2-MR-CAR T cells were more effective at trafficking to and lysing the tumor cells compared to the HER2-CAR T cells.

Finally, we looked into whether the tumor-infiltrating CAR T cells still had enhanced capacity to migrate, proliferate and lyse tumor cells. HER2-positive A549 tumor cells were co-cultured for 24 hours with the sorted HER2-MR-CAR T cells, HER2-CAR T cells, or non-transduced T cells at a ratio of 5:1. The culture media were then collected for quantitating IFN-γ and LDH as indicators of T cell’s activation and cytotoxicity, respectively **(Figure 6E-F)**. The results showed that HER2-MR-CAR T cells exhibited heightened activation and cytotoxicity against A549 cell, compared to HER2-CAR T cells and non-transduced T cells. Additionally, the sorted HER2-MR-CAR T cells showed significantly enhanced migration capability compared to HER2-CAR T cells and non-transduced T cells when assessed using a transwell migration assay **(Figure 6G)**. Moreover, the proliferation capability of HER2-MR-CAR T cells was significantly stronger compared to that of HER2-CAR T cells and non-transduced T cells as assessed by staining with CFSE and analyzing their proliferation using flow analysis **(Figure 6H)**. In conclusion, these findings suggest that HER2-MR-CAR T cells retained their capability to proliferate, migrate, and lyse tumor cells in the tumor microenvironment.

## Discussion

We here report on a CAR T cell therapy that metabolically refuels (MR) T cells with inosine using a helper vector that encodes CD26 and ADA1 fused to anti-CD3 scFv. Our MR-CAR T cell therapy has the potential to address the challenges of current CAR T cell therapy in treating solid tumors by overcoming the immunosuppressive mechanisms present in the tumor microenvironment. To achieve this, our MR-CAR T cell strategy implements three innovative mechanisms: i) ADA1 mediates the conversion of adenosine to inosine, which overcomes adenosine-mediated immunosuppression of CAR T cells; ii) inosine selectively promotes CAR T cell proliferation in the nutrient-deprived tumor microenvironment without feeding tumor cells; and iii) the overexpression of CD26 provides optimal co-stimulatory signals to CAR T cells, improving their mobility and enhancing anti-tumor effects **(Figure 7)**.

**Figure 7.**
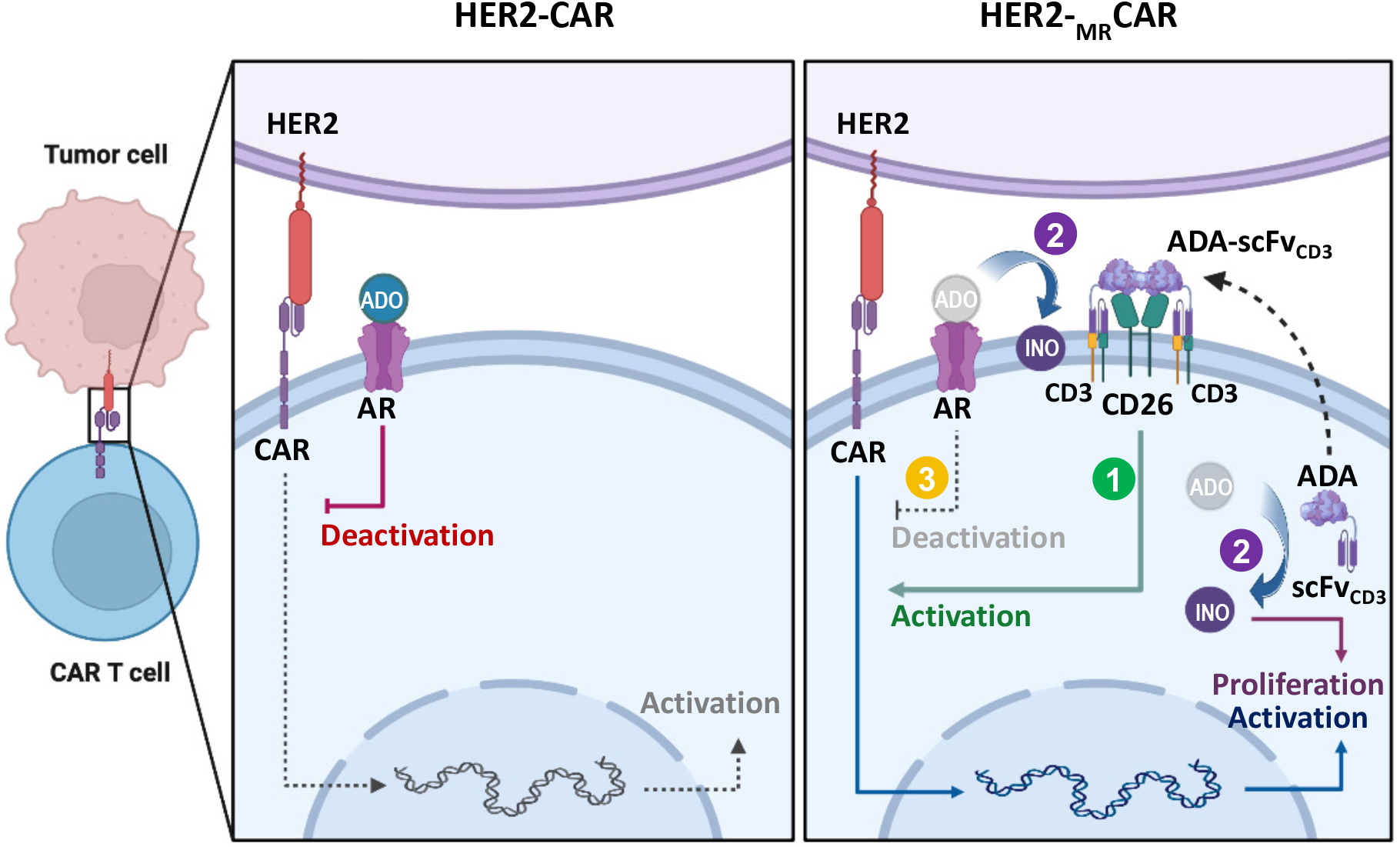
Rational for MR-CAR T cell therapy. MR-CAR T cells express a tumor-specific CAR and CD26/ADA1.CD3scFv encoded by the MR-vector. CD26 and ADA1.CD3scFv enhances the anti-tumor activity of CAR-T through different mechanisms: (1) CD26 mediates CAR T cell trafficking and activation; (2) ADA1 provides both intracellular and extracellular INO as carbon source for CAR T cell proliferation; and (3) ADA1 releases the ADO-mediated immunosuppression.

First, adenosine signaling is an important immuno-metabolic checkpoint in tumors^23,55^. Ecto-nucelotidases, such as CD39, CD73, CD38, CD203a, ALP, and PAP can generate adenosine from ATP or NAD+ that accumulate in the tumor microenvironment^20,56–60^. Adenosine enables tumor cells to evade immune surveillance by suppressing the function of various protective immune cells, such as T cells, DCs, NK cells, macrophages, and neutrophils, while promoting the activity of immunosuppressive cells, such as myeloid-derived suppressor cells (MDSCs) and regulatory T cells (Tregs)^33,58,61–68^. Adenosine also activates cancer-associated fibroblasts and induces the formation of new blood vessels^64,69^. Many drugs have been discovered, including small molecules or monoclonal antibodies targeting CD73 and CD39 to limit adenosine production or A2AR and A2BR to inhibit adenosine binding to immune cells^55,62^. Their anti-tumor effectiveness has been shown in preclinical investigations, both by themselves and in conjunction with other immunotherapies such immune checkpoint inhibitors and adoptive cell transfer. However, early-phase clinical studies have failed to demonstrate adequate anti-tumor effects. Existing adenosine-targeted therapies only shut down one particular adenosine pathway while leaving the others unaffected whereas multiple ecto-nucleotidases contribute to extracellular adenosine production, and adenosine binds to multiple receptors, such as A2AR, A2BR, and A3R, to suppress antitumor immunity. ADA1 offers an alternate strategy to current adenosine-target specific medicines for combating adenosine-mediated immunosuppression by converting it to inosine.

Second, we have shown in earlier studies that inosine can serve as an alternative energy source to support T cell proliferation and function in the absence of glucose^19^. Cancer cells with high glycolytic activity may deplete glucose, leading to a shortage of glucose in T cells. To enhance T-cell based therapies, supplementing T cells with inosine has been proposed. Nevertheless, tumor cells also have the ability to utilize inosine. Hence, it is imperative to develop a strategy to specifically deliver inosine to CAR T cells while avoiding tumor cells. In this regard, we overexpressed ADA1 in the cytoplasm of CAR T cells, which increased the generation of intracellular inosine, thereby boosting the growth of CAR T cells. Furthermore, upon conditional release in the tumor environment, the anti-CD3 scFv will engage ADA1 and the CAR T cells. The ecto-ADA1 on CAR T cells will convert adenosine in the T cell surroundings to inosine, thus overcoming adenosine-mediated immunosuppression and increasing inosine concentration around CAR T cells. Moreover, the CD3-scFv will bind to endogenous CD3+ T cells, resulting in bystander effects.

We explored different options for using ADA1 in CAR T cell therapy, including overexpressing cytoplasmic ADA1, secreted ADA1, and membrane-bound ADA1. However, we found that overexpressing secreted ADA1 was not feasible since its secretion is not inducible, even with the addition of an IL2 signal peptide. Qu et al. reported only a marginal increase in ADA1 secretion upon CAR T cell induction in response to IL2 signal peptide, which could be due to the stress condition present in their cell culture system^34^. Overexpressing the wild-type cytoplasmic ADA1 in CAR T cells could also be problematic since its conditional secretion could provide inosine as an energy source to tumor cells. While ADA1 is a ubiquitously expressed cytoplasmic protein, it can be secreted in response to stress stimuli such as hypoxia and inflammation, and the secretion is mediated by a non-classical pathway that does not require a signal peptide. This is similar to a subset of proteins, including IL-1α, IL1-β, IL-18, IL-33, IL-36α, IL-37, and IL-38, which lack a signal peptide and do not follow the classical endoplasmic reticulum-to-Golgi pathway of secretion^70,71^. Instead, these proteins were likely transport out of the cell via exocytosis of preterminal endocytic vesicles during inflammatory or stress conditions. Indeed, elevated ADA1 expression in human tumors has been detected and linked to poor survival outcome^25^. Furthermore, our data demonstrated that permanent membrane-bound ADA1 can induce autocrine activation in a tumor antigen-independent manner, likely due to CD26 engagement. Hence, anchoring ADA1to T cells through CD3scFv and CD26 is the most efficient way to use inosine in CAR T cells. Cytoplasmic ADA1 converts intracellular adenosine to inosine, and upon secretion, produces inosine in a CAR T cell membrane-proximal manner, which avoids feeding tumor cells.

Third, our strategy overexpresses CD26 on the surface of CAR T cells as the CD26 expression level on the unmodified CAR T cells is relatively low and can be further downregulated by TGF-ß1, a cytokine found in significant quantities in various tumor tissues^72–78^. Thus, we develop a strategy to overexpress CD26 on the surface of CAR T cells, which sustains co-stimulation and membrane proximal ADA1 capture even in TGF-ß1-rich environment and provides a survival advantage for these engineered therapeutic cells.

To our knowledge, this study is the first to explore the use of a T cell-anchoring ADA1 and overexpressed CD26 in CAR T cell therapy. Data herein suggest that MR-CAR strategy has the potential to overcome the limitations of current CAR T cell therapy for solid tumors by providing inosine as an alternative carbon source and overcoming adenosine-mediated immunosuppression. In preclinical NSCLC models, HER2-specific MR-CAR T cells exhibited enhanced anti-tumor activity. HER2 is overexpressed in certain types of cancers, including non-small cell lung cancer (NSCLC), breast, gastric, and ovarian cancers. Although two HER2-targeted ADCs (DS-8201 and T-DM1) have been approved by FDA^79–81^, they have serious toxicities, such as grade 3+ adverse events, treatment discontinuation, interstitial lung disease or pneumonitis, and even deaths. Additionally, HER2-ADCs have shown limited efficacy in some patients with advanced HER2-positive cancers, likely due to a lack of effective internalization and intracellular delivery of the drug payload, as well as the restricted ability of these ADCs to cross the intact blood-brain barrier (BBB). Resistance to HER2-ADCs can develop over time, as the cancer cells evolve to evade the effects of the drug payload. This can limit the long-term effectiveness of HER2-ADC and result in disease progression. Thus, HER2-specific MR-CAR T cell strategy may offer a safter and more effective alternative for patients with HER2-positive cancers^43,45,82^. Additionally, our MR-CAR T cell strategy may be applicable to a broad range of T cell-based therapies currently under investigation, including autologous CAR T cell therapy, allogenic CAR T cell therapy, TIL therapy, and in vivo CAR therapies.

## Disclosure of Potential Conflicts of Interest

X.S., K.S., and A.S., have equity interests in Cellula Biopharma, Inc., the company that intends to commercialize the technology discussed in this work.

## Author Contributions

X.S., Y.H., and A.S. designed experiments, analyzed data, created the figures, and wrote/edited the manuscript; K.S., executed experiments and wrote/edited the manuscript; S.M. executed experiments; M.H., R.W., and A.H. provided intellectual feedback and edited the manuscript. All authors critically read and approved the manuscript.

## Acknowledgments

This study was supported by Department of Defense Lung Cancer Research Program Idea Award W81XWH2210701 and Texas A&M Translational Investment Fund to X Song. We thank Dr. Arijita Sarkar (University of Southern California) and Dr. Saikat Chowdhury (The UT MD Anderson Cancer Center) for helping to analyze the RNAseq data.

## Data availability

All data are available from authors upon reasonable request.

---

**sFigure 1.**
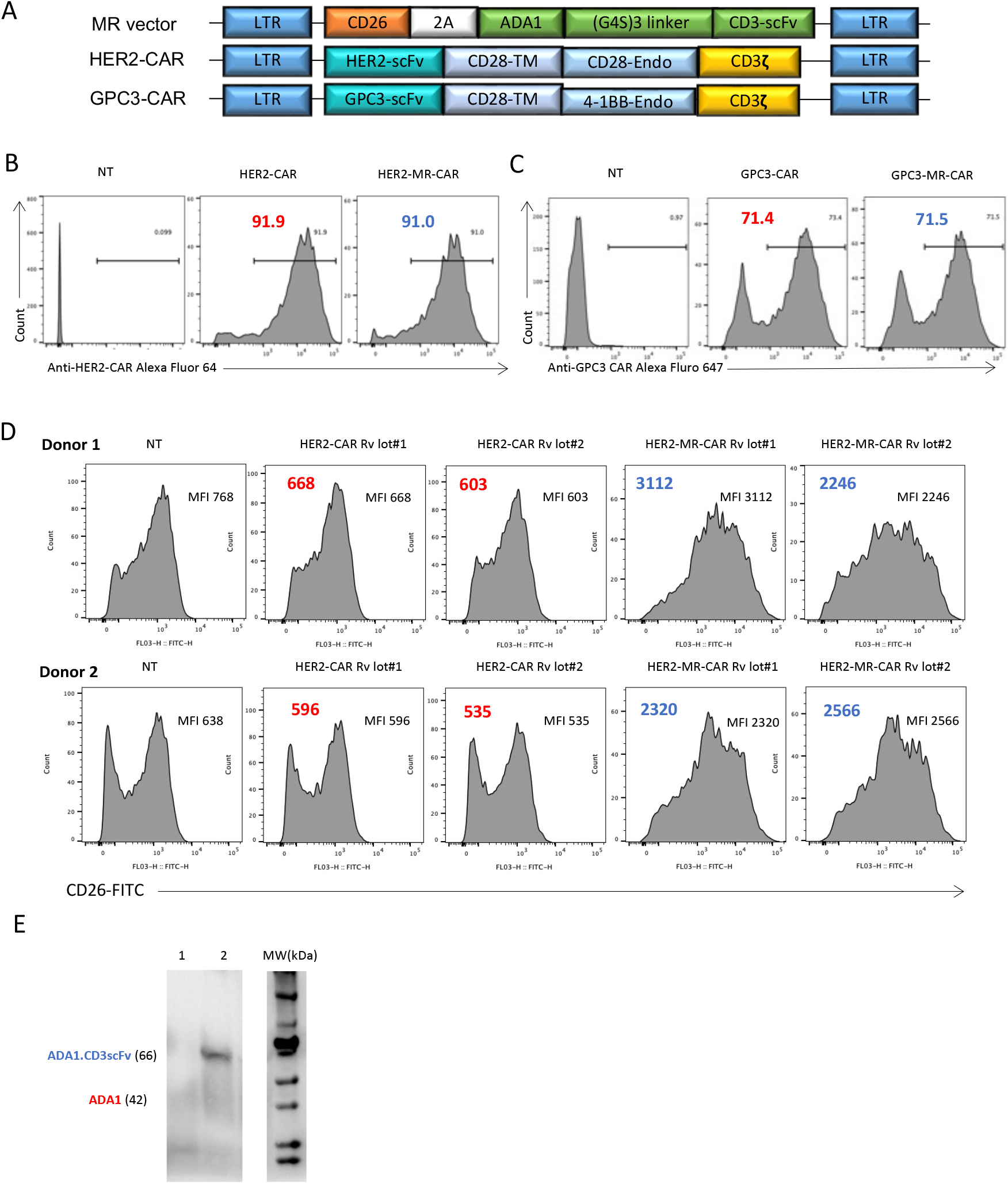
Characteristic of MR-CAR T cells. **(A)** Schematic representation of MR vector encoding CD26 and ADA1.CD3scFv, HER2-CAR vector encoding HER2-specific scFv fused with CD28-TM, CD28-endodomain, and CD3ζ, and GPC3-CAR vector encoding GPC3-specific scFv fused with CD28-TM, 4-1BB-endodomain, and CD3ζ. **(B)** Flow analysis was performed to detect HER2-specific CAR expression using PE-conjugated human HER2 protein. **(C)** Flow analysis was performed to detect GPC3-specific CAR expression using PE-conjugated anti-Fab Ab. **(D)** CD26 expression on HER2-MR-CAR T cells, HER2-CAR T cells, or non-transduced T cells (NT) was detected by flow analysis using anti-CD26 mAb. **(E)** Western blotting analysis was performed to detect ADA1.CD3scFv. Cell lysates of HER2-CAR T cells (lane 1) and HER2-MR-CAR T cells (lane 2) were analyzed using anti-ADA1 antibody.

**sFigure 2.**
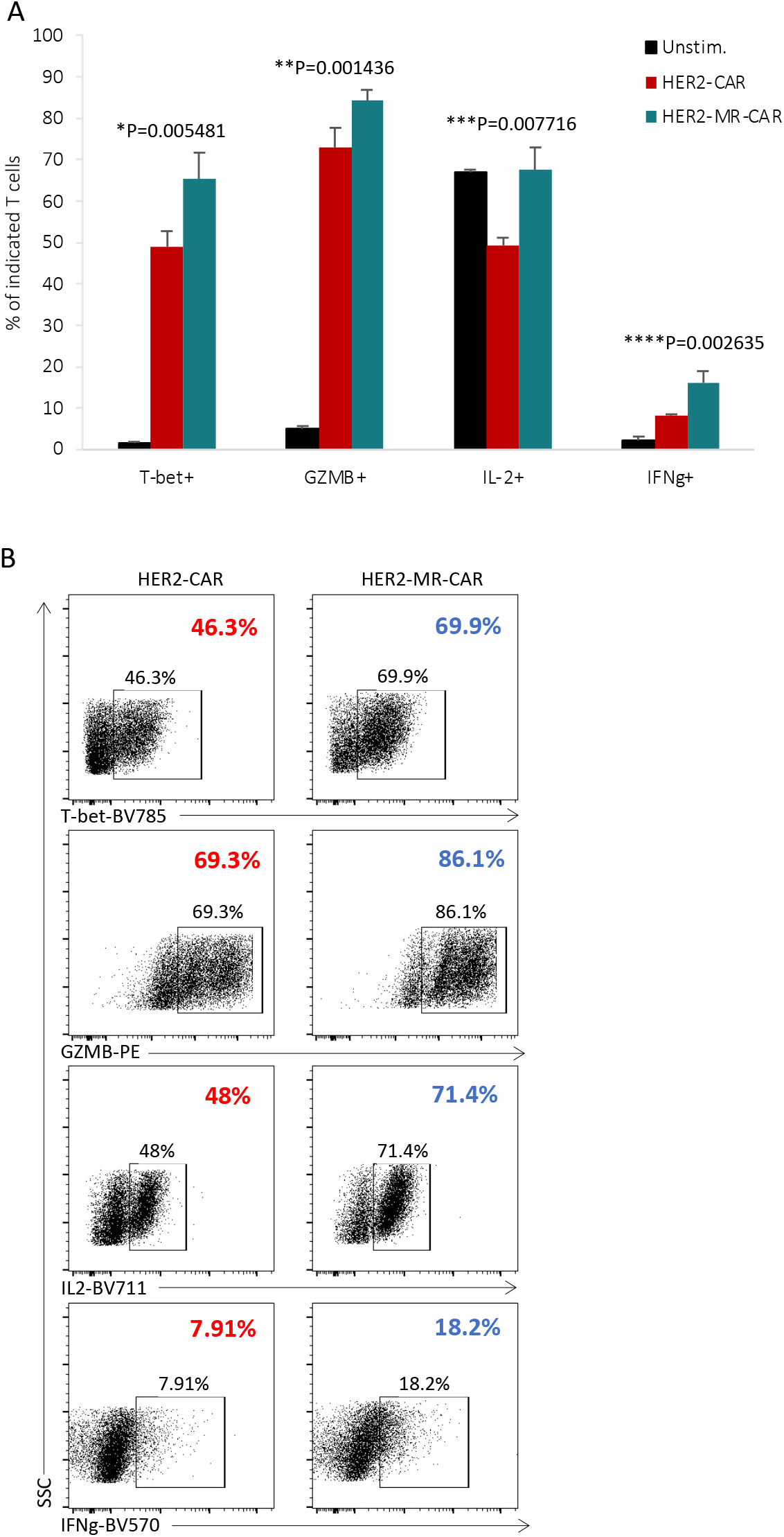
MR-CAR T cells exhibit higher expression of T-bet, granzyme B, IFN-γ, and IL-2 following CD3 antibody stimulation. **(A)** HER2-MR-CAR T cells and HER2-CAR T cells were re-stimulated with a CD3-coated plate overnight, and subsequently analyzed by flow cytometry to assess expression of T-bet, granzyme B, IFN-γ, and IL-2. The figure indicates the P values. **(B)** Representative data is shown.

**sFigure 3.**
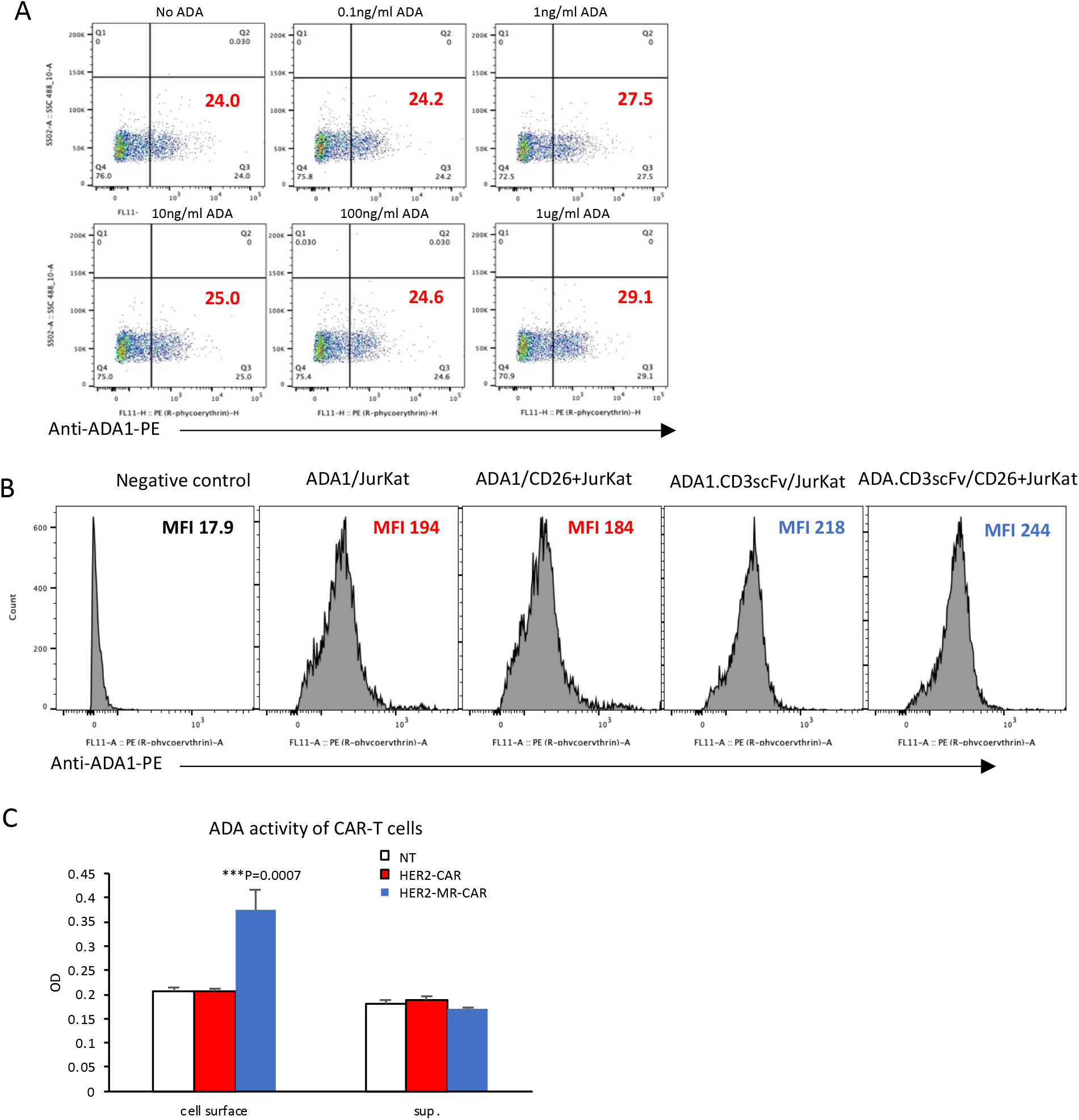
CD3scFv facilitates binding of ADA1.CD3scFv and T cells. **(A)** CD26-positive Jurkat T cells were cultured with ADA1 protein at different concentrations for 24 hours. The binding of ADA1 to Jurkat T cells were detected by flow analysis using anti-ADA1 antibody staining. **(B)** CD26-negative or CD26-positive Jurkat T cells were transduced with retroviral vectors encoding ADA1 or ADA1.CD3scFv. The binding of ADA1 to Jurkat T cells were detected by flow analysis using anti-ADA1 antibody staining. **(C)** HER2-MR-CAR T cells and HER2-CAR T cells were generated from human PBMC activated by anti-CD3/CD28 antibodies and transduced with HER2-CAR vector with or without MR vector. After 48 hours of culture, the cells or culture medium were analyzed for ADA activity following the manufacturer’s protocol. *P=0.0007 for HER2-MR-CAR verse HER2-CAR.

**sFigure 4.**
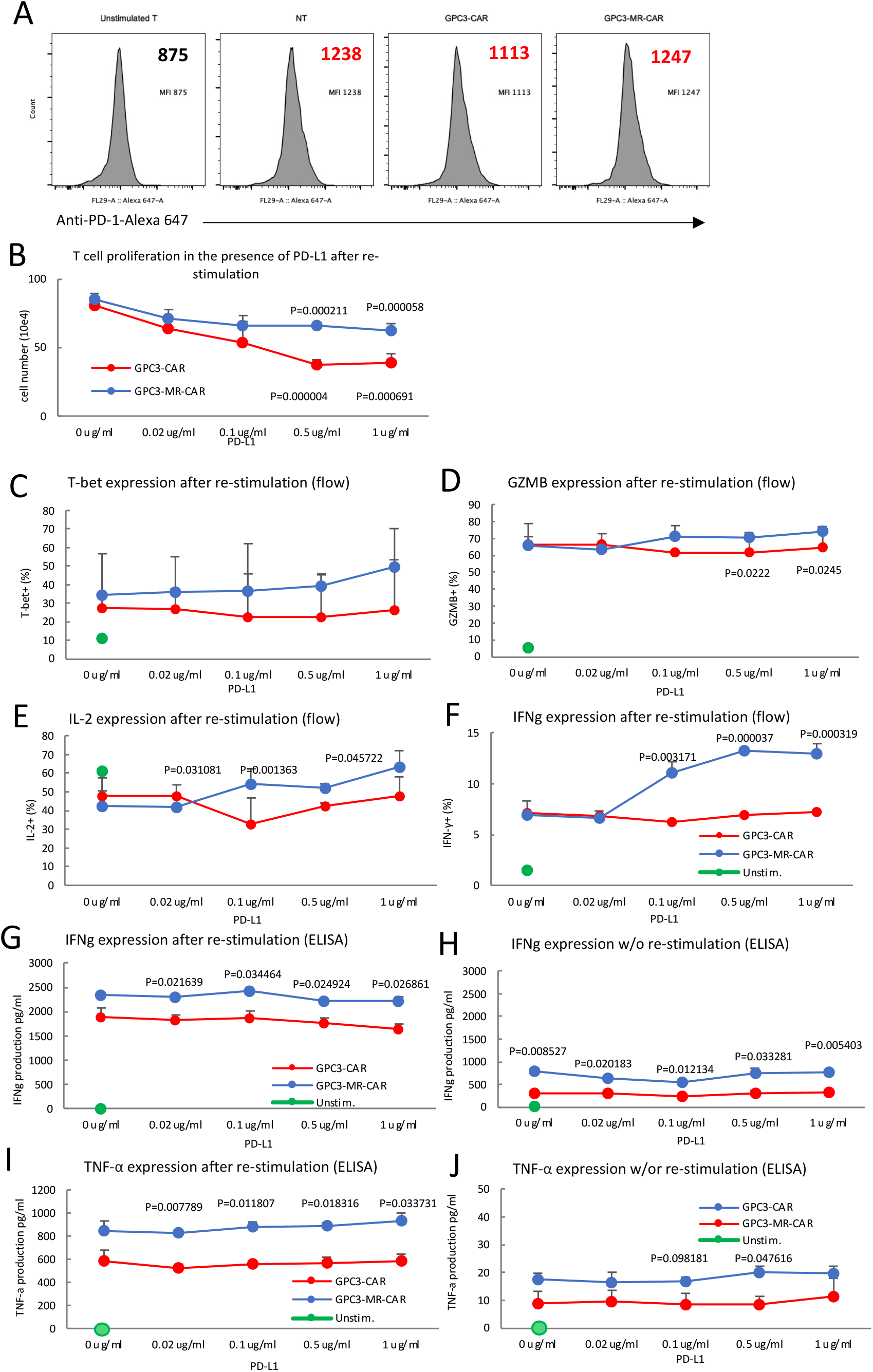
MR-CAR T cells exhibit enhanced growth and effector functions compared to CAR T cells under PD-L1 inhibition. **(A)** Flow analysis of PD-1 expression was conducted on GPC3-MR-CAR T cells, GPC3-CAR T cells, non-transduced T cells, and unstimulated T cells. **(B-J)** 1×10^6^ of GPC3-MR-CAR T cells and GPC3-CAR T cells were cultured overnight in the presence of PD-L1 at the indicated concentration. Cell count was determined using blue stain (B). The cells were collected and analyzed for T-bet (C), granzyme B (D), IL-2 (E), and IFN-γ (F) expression using flow analysis. ELISA was used to measure IFN-γ with re-stimulation (G) or without re-stimulation (H) and TNF-α with re-stimulation (I) or without re-stimulation (J). The figure indicates the P values.

**sFigure 5.**
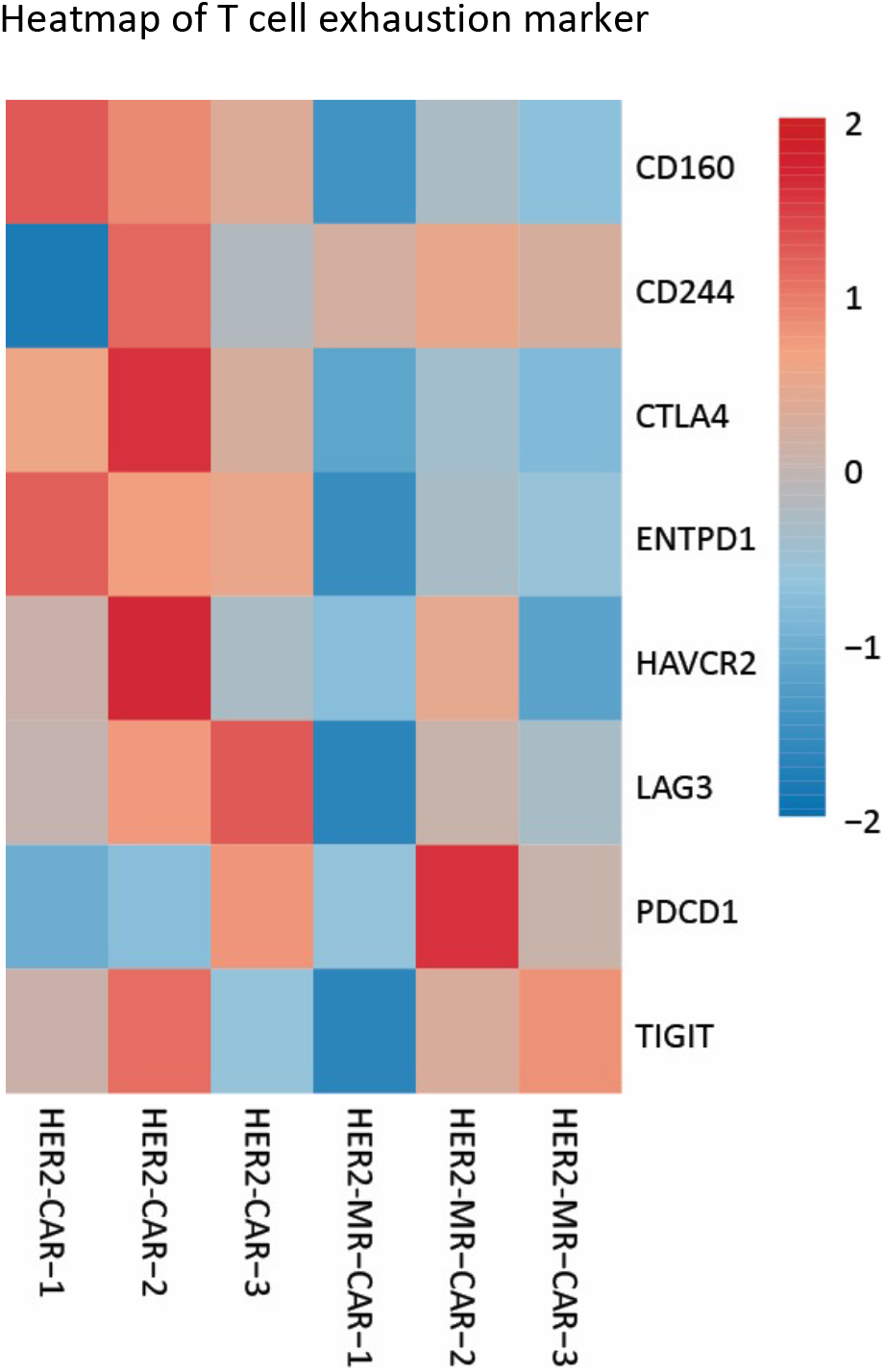
Heatmap showing T cell exhaustion marker genes expression. The heatmap shows the expression levels of T cell exhaustion marker genes in two groups, represented on the Y-axis. The expression levels were converted to Z-scores to enable comparison between the different cytokine genes. Triplicate samples (N=3) were used for each group and are represented on the X-axis. The heatmap provides a visual representation of the relative expression levels of T cell exhaustion marker genes in the two groups, with warmer colors indicating higher expression levels and cooler colors indicating lower expression levels.

**sFigure 6.**
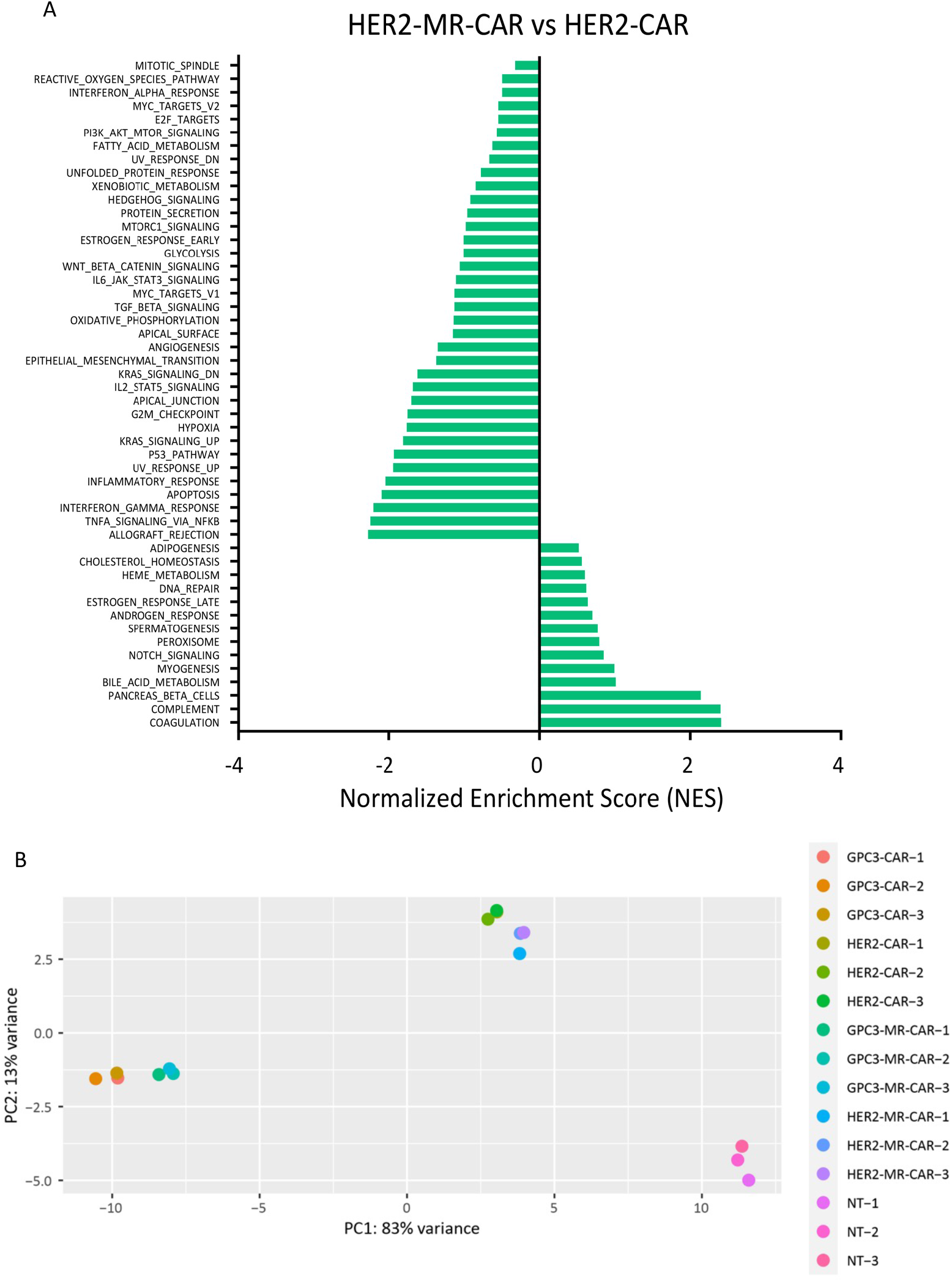
Hallmark GSEA of RNA-seq data performed on HER2-MER-CAR T cells vs HER2-CAR T cells. (**A)** Hallmark Gene Set Enrichment Analysis (GSEA) was performed on RNA-seq data from HER2-MER-CAR T cells and HER2-CAR T cells using triplicate samples for both groups. The figure shows the results based on the normalized enrichment score (NES), which measures the degree of pathway enrichment in the HER2-MR-CAR T cells compared to the HER2-CAR T cells. Positive NES values indicate pathway upregulation in the HER2-MR-CAR T cells, while negative NES values indicate downregulation. The figure provides a visual representation of the pathways that are significantly enriched in the HER2-MR-CAR T cells compared to the HER2-CAR T cells. (**B)** Principal Component 1 (PC1) explains 83% of the variance observed in the Hallmark Gene Set Enrichment Analysis (GSEA) results.

**sFigure 7.**
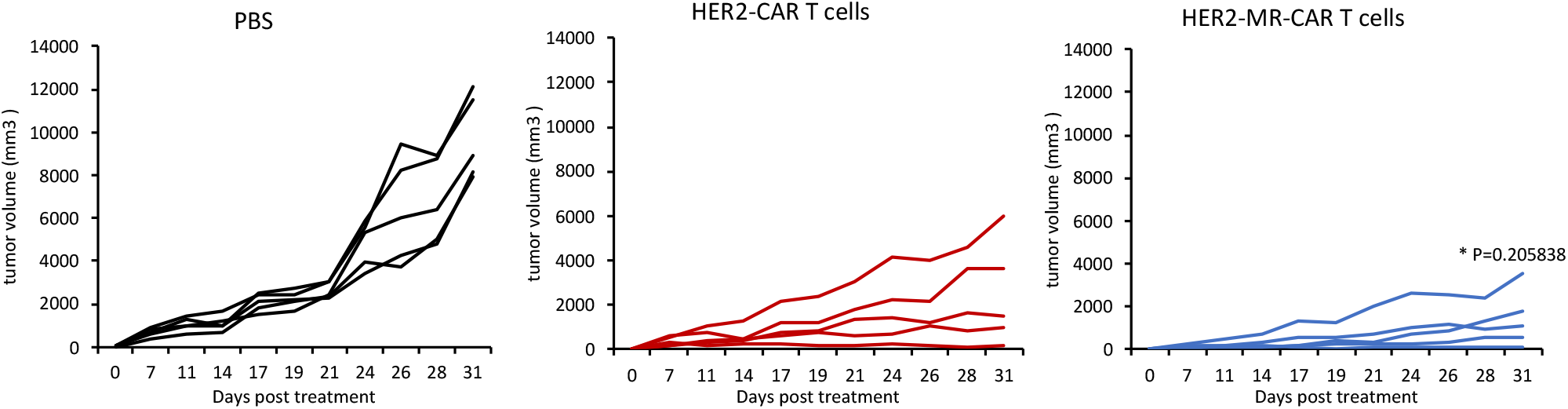
HER2-specific MR-CAR T cells demonstrated anti-tumor activity in a Calu3 xenograft mouse model. A murine melanoma xenograft model was established in NSG mice by subcutaneous inoculation of 2 × 10^6^ Calu3 tumor cells on the right flank. When the average tumor size reached 4-6 mm in diameter, experimental mice were treated with a single dose of either 2 × 10^6^ HER2-MR-CAR T cells, 2 × 10^6^ HER2-CAR T cells, or PBS. Tumor size and mouse body weight were monitored every two to three days. Data represent mean ± s.d. (n = 5). The results showed that treatment with HER2-MR-CAR T cells resulted in a trend towards reduced tumor growth compared to treatment with HER2-CAR T cells (P=0.205838, HER2-MR-CAR T cells vs HER2-CAR T cells on day 31).

## Notes

### Competing Interest Statement

X.S., A.S., and Y.H. are the inventors of the technology discussed in this work, and Texas A&M University has ownership of the technology and has filed a patent application for it.

